# Neutrophil transcriptome diverges into two discrete trajectories in a murine model of severe *Streptococcus pneumoniae* pneumonia

**DOI:** 10.1101/2024.10.29.620672

**Authors:** Riley M.F. Pihl, Salam H. Alabdullatif, Bradley E. Hiller, Elise M.R. Armstrong, Kevyn R. Martins, Ernest L. Dimbo, Yewoo Lee, Joshua D. Campbell, Adam C. Gower, Joseph P. Mizgerd, Lee J. Quinton, Katrina E. Traber

## Abstract

Neutrophils were once considered a homogenous population of transcriptionally static, pathogen-killing cells, however, recent models have demonstrated neutrophil functional and transcriptional plasticity. We performed transcriptomic analyses in a murine model of pneumococcal pneumonia to investigate neutrophil plasticity and demonstrate that neutrophils are highly dynamic, leading to three distinct alveolar neutrophil populations – one immature (early bronchoalveolar lavage neutrophils [BALN]) and two mature (late BALN). Early BALNs produce high levels of inflammatory cytokine transcripts, maturing into late BALNs, including a pro-degranulation and phagocytosis population (late-degranulating BALN) or a population specializing in translation machinery and inflammatory cytokine production (late-cytokine producing BALN). Neutrophil metabolism is also regulated in a stepwise manner – tricarboxylic acid (TCA) cycle and respiratory electron transport chain (ETC) genes are downregulated as neutrophils migrate from the vasculature to the interstitium, lipid and carbohydrate metabolism genes are downregulated during migration from interstitium to the airspace. These transitions may be regulated by aspects of the integrated stress response (ISR), as key regulators including *Eif2ak2* are upregulated in interstitial neutrophils. Overall, we demonstrate that pneumonic neutrophils are transcriptionally plastic, developing through two distinct transcriptional phenotypes in the airspace, and are metabolically and transcriptionally rewired with potential points of regulation occurring in the interstitial space.

## Introduction

Despite pathogen-specific vaccines and antibiotics, including those effective against the most common bacterial cause of community-acquired pneumonia, *S. pneumoniae*, respiratory infections remain a leading cause of global deaths (1). Immune dysregulation, coinciding with unregulated inflammatory cytokine production and/or exuberant innate immune effector function often can promote tissue damage, negatively affecting patient outcomes (2, 3). Targeting immune dysfunction is the primary goal of host-directed pneumonia therapeutics, yet our understanding of this complex problem remains limited (3, 4).

Neutrophils are an integral component of the innate immune response in pneumonia and can drive immune dysfunction. Neutrophils have been reported to drive tissue damage in acute respiratory distress syndrome (ARDS) via inflammatory signaling and effector functions, but somewhat paradoxically low neutrophil to lymphocyte ratios track with progression of ARDS (5, 6). Canonically, neutrophils have been characterized as short-lived innate immune effector cells responsible for pathogen clearance. They migrate to the site of infection, and kill pathogens through multiple mechanisms; phagocytosis, degranulation releasing reactive oxygen species (ROS) and enzymes including myeloperoxidase (MPO) and gelatinase B (MMP9), and/or production of neutrophil extracellular traps (NETs) (7). Work in recent years has shown that neutrophils can have numerous additional functions beyond pathogen-killing, including T and B cell regulation (8, 9), roles in cancer propagation or elimination (10), and resolution of inflammation (11). Further, neutrophils are increasingly proving to be transcriptionally dynamic, and highly adaptive to the tissue environment where they are located. Neutrophils isolated from different tissues and in different disease conditions have drastically different transcriptional and functional states (12–15), indicating that their plasticity may result from location-specific environmental cues, including those within and en route to infected airspaces.

As central players in early pneumonia responses, neutrophils are an attractive target, and their modulation may to help manage infection while limiting secondary lung injury (16–18). While a new picture of neutrophils as transcriptionally dynamic and plastic first-responders is emerging, the neutrophil transcriptional rewiring during migration into the lung during pneumonia as well as how such dramatic transcriptional changes are regulated is poorly understood. Therefore, the goal of this work is to determine lung-specific transcriptional programs in neutrophils and identify possible mechanisms of regulation.

## Results

### Neutrophil transcriptome remodeling in the alveolar space

To begin our investigation of the neutrophil transcriptome during pneumonia, we used gene expression microarrays to profile RNA obtained from peripheral blood and BALF neutrophils isolated from mice infected with *S. pneumoniae* serotype 3 (SP3) or saline control for 24 hours. Neutrophils were isolated by fluorescence activated cell sorting (FACS) of peripheral blood from infected and uninfected mice, and BALF from infected mice (**Figure 1A**; **Supplemental Figure 1A-B**). Differentially expressed genes (DEGs) (one-way ANOVA, FDR *q* < 0.05) were identified and used to perform hierarchical clustering (**Supplemental Figure 1C**). Interestingly, thousands of genes were differentially expressed (FDR *q* < 0.1) between alveolar and peripheral blood neutrophils in infected animals (**Figure 1B**), but there were few significant changes in peripheral blood neutrophils between uninfected and infected mice (**Figure 1C**). This suggests that significant transcriptomic changes occur during migration from the circulation to the alveolar space but not in response to infection itself. The top 20 upregulated and downregulated genes between peripheral and alveolar neutrophils in infected mice are listed (Supplemental **Table 1,** full list in **Supplemental Table 2**). Top upregulated genes in alveolar neutrophils include inflammatory cytokines such as *Il1a* and *Tnf*, as well as immune cell chemokines such as *Ccl3, Cxcl10,* and *Ccl4*. To validate results from the array, qRT-PCR from a separate set of experiments was used to directly quantify RNA of selected genes in neutrophils isolated from peripheral blood or alveolar space of mice infected with SP3 for 24h (**Supplemental Figure 1D-E**).

**Figure 1.**
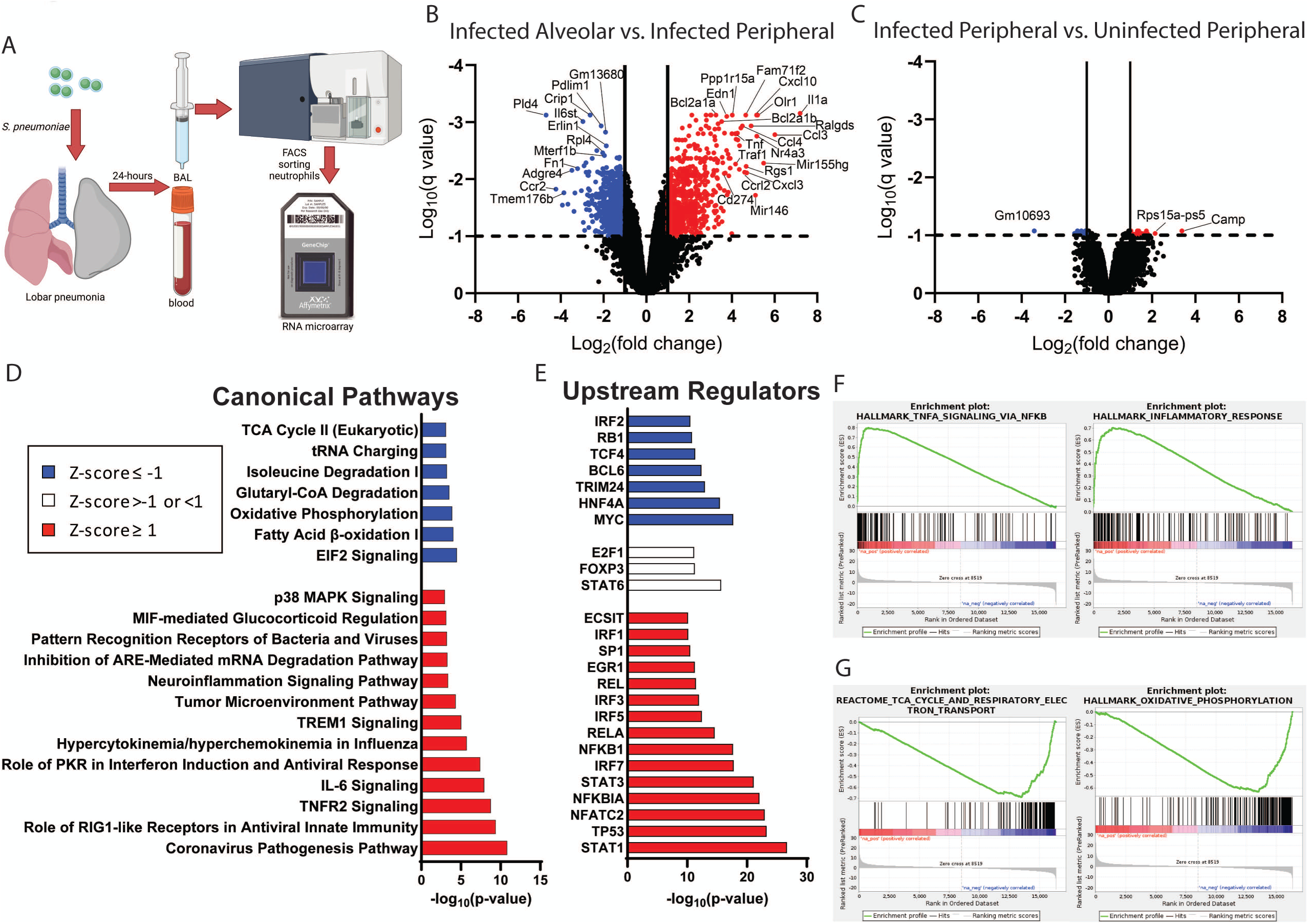
Neutrophil transcriptome remodeling in the alveolar space. (A) Schematic of experimental design. Mice were infected with ∼ 1.0×10^6^ CFU *S. pneumoniae* (SP3) or saline control for 24 hours. Neutrophils were collected from BAL of infected mice or from the blood of infected or uninfected mice, isolated via FACS, and profiled by Affymetrix gene expression microarray. Peripheral and alveolar neutrophils of infected mice were isolated from the same animal. (B-C) Volcano plot of all genes after expression filtering, log_10_(q value) vs log_2_(fold change) with selected DEG labeled for (B) infected-alveolar vs. –peripheral and (C) infected peripheral vs. uninfected peripheral. Blue and red dots represent genes down-or up-regulated, respectively, more than 2-fold (vertical solid lines), and FDR *q* < 0.1 (horizontal dashed lines). (D) Pathway activation and (E) upstream regulators as predicted by Ingenuity Pathway Analysis (IPA). Bars colored based on Z-score of predicted change [blue < –1, white 0 and red >1]. Significance determined by uncorrected *p* value (right). (F-G) Example Gene Set Enrichment Analysis (GSEA) plots of pathways with coordinately (F) upregulated or (G) downregulated genes. N=3 female mice.

To understand the biologic processes represented by the gene changes between peripheral blood and alveolar neutrophils, we performed several *in silico* analyses. Using Ingenuity Pathway Analysis (IPA, Qiagen, (19)), we identified canonical pathways and predicted upstream regulators from our dataset (**Figure 1D-E**). Top upregulated pathways involve pathogen sensing (PRR recognition, RIG1-like receptors) and pro-inflammatory responses to pathogen (IL-6 signaling, TNFR2 signaling). Downregulated pathways affect tRNA charging, metabolism (oxidative phosphorylation, Fatty Acid β-oxidation, TCA Cycle), and regulators of stress responses (EIF2 signaling). Activated upstream regulators included those involved in NFκB signaling (*Nfkb1, RelA*) and interferon signaling (*Stat1, Irf3, Ifr5, Irf7*), while those repressed or downregulated include those involved in cell cycle (*Rb1* and *Myc*) and chromatin (*Irf2* and *Trim24*). We also performed Gene Set Enrichment Analysis (GSEA, (20, 21)), ranking all genes interrogated by the microarray by the moderated *t* statistic for infected alveolar vs peripheral neutrophils, and using gene sets from the Molecular Signatures Database (MSigDB). Lists of the top 20 gene sets coordinately up-or down-regulated were generated (**Supplemental Table 2**; full list in **Supplemental Table 3**), and **Figures 1F-G** depict example GSEA plots. Confirming the findings from IPA, we again see a strong enrichment in inflammatory pathways involving NFκB, interferon, and STAT signaling, and a downregulation of cellular machinery, metabolism, and cell cycle pathways.

Overall, these data suggest that as neutrophils migrate from the circulation, inflammatory genes are dramatically upregulated, and genes associated with cellular machinery are downregulated. This is consistent with the dogma that neutrophils travel to the site of infection to act as inflammatory effector cells.

### Neutrophil transcription is highly distinct between blood and BAL and dynamic within blood or BAL

Sequencing depth afforded by the bulk transcriptomics approach illustrated in **Figure 1** provided essential evidence of neutrophil reprograming between the circulating and intra-pulmonary compartments. We next sought to determine whether transcriptomic differences are a result of global changes or specific subsets. Single cell RNA sequencing (scRNA-seq) was performed on neutrophils isolated from blood and BALF isolated from mice infected with SP3 for 24 hours. To preserve neutrophil cellular integrity and improve RNA yield, neutrophils were enriched using negative-selection magnetic beads and profiled with the 10X platform (**Figure 2A**). Initial clustering with all cells was performed with Celda (22) (**Supplemental Figure 2A**). Because neutrophils were enriched rather than FACS sorted, we anticipated ∼5-10% contamination by non-neutrophils. Using markers for other cell types from a previously published lung single cell atlas (LungMap Cell Cards) (23), we identified and excluded non-neutrophils from further analyses (**Supplemental Figure 2A-B**). After re-clustering the remaining neutrophils, eight clusters were identified; blood neutrophil were primarily in clusters 1 (C1) and 2 (C2), and BAL neutrophils (BALN) in clusters 3-8 (C3-C8) (**Figure 2B**). In agreement with our microarray data from **Figure 1**, gene expression in blood neutrophils was drastically different from BALNs as these two groups separate completely in the UMAP plot. We then examined transcripts that were top DEGs between each cluster (**Figure 3C**). *Ly6g,* and *Cd177* mRNA are expressed almost exclusively by blood neutrophils (**Figure 2C-D**). *Ly6g* is a common marker for neutrophil maturation, and *Cd177* is a marker of activation and maturation in neutrophils (24–26). Their near exclusive expression by blood neutrophils suggests that a critical threshold of neutrophil transcriptional maturation occurs in the blood, and these genes are no longer necessary once neutrophils have migrated to the infected lung. Amongst the two blood clusters, the immune regulator *Cd244* and *Hdac4* are preferentially transcribed by C1, while the markers of maturation and activation *Ly6G, Ngp,* and *CD177* are transcribed by C2 neutrophils (**Figure 2C-D**). HDAC4 is a key histone deacetylase that regulates numerous transcriptional pathways involved in growth and development (27). The differential expression of these genes suggest that neutrophil maturation occurs in two distinct phases in the blood, one where neutrophil effector and activation markers (*Ly6g, Ngp,* and *Cd177*) are transcribed, and another defined by regulatory factors (*Cd244* and *Hdac4*). Among the BALN, expression of markers of inflammation, chemokines, and pathogen response including *Cxcl10, Il1a, Tnf, Icam1, Cxcr4, F10, Osm, Cxcl3,* and *Ifit3* are upregulated in BALN compared to blood and stratifies the different BALN clusters, as discussed further below (**Figure 2C-D**).

**Figure 2.**
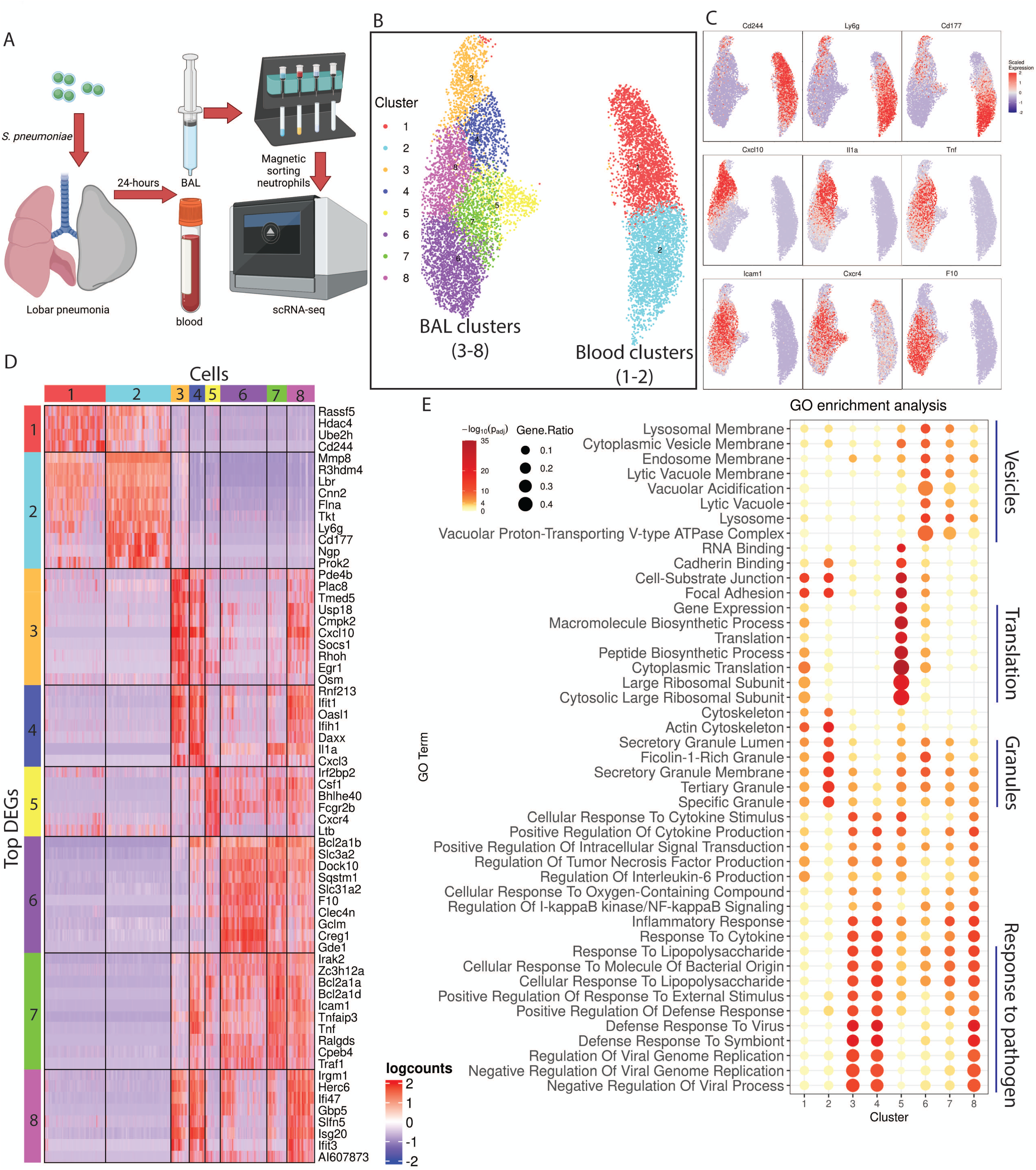
Neutrophil transcription is highly distinct between blood and BAL and dynamic within blood or BAL. (A) Experimental schematic. Neutrophils isolated from the blood and BAL of SP3-infected animals via negative selection magnetic bead enrichment followed by scRNA-seq. (B) Neutrophils were clustered via the Celda algorithm and plotted on a UMAP. (C) Scaled expression of select genes plotted on a UMAP. (D) The top 10 DEGs from each cluster (overlap in top genes results in consolidated list) in a heatmap where rows and columns correspond to genes and cells, respectively; columns further arranged by Celda clusters; colors are scaled by row, with blue and red indicating values that are at least 2 standard deviations below or above the mean (white), respectively. (E) Gene Ontology (GO) enrichment scoring was performed on DEGs (significance p < 0.05 and log_2_(fold change)>0.25) from each cluster and the top 10 enriched pathways for each cluster are shown, note some pathways are shared by multiple clusters. Pathways and clusters are ordered hierarchically by gene ratio and both the gene ratio and adjusted P-value for each pathway and cluster are shown. N=2 male mice.

**Figure 3.**
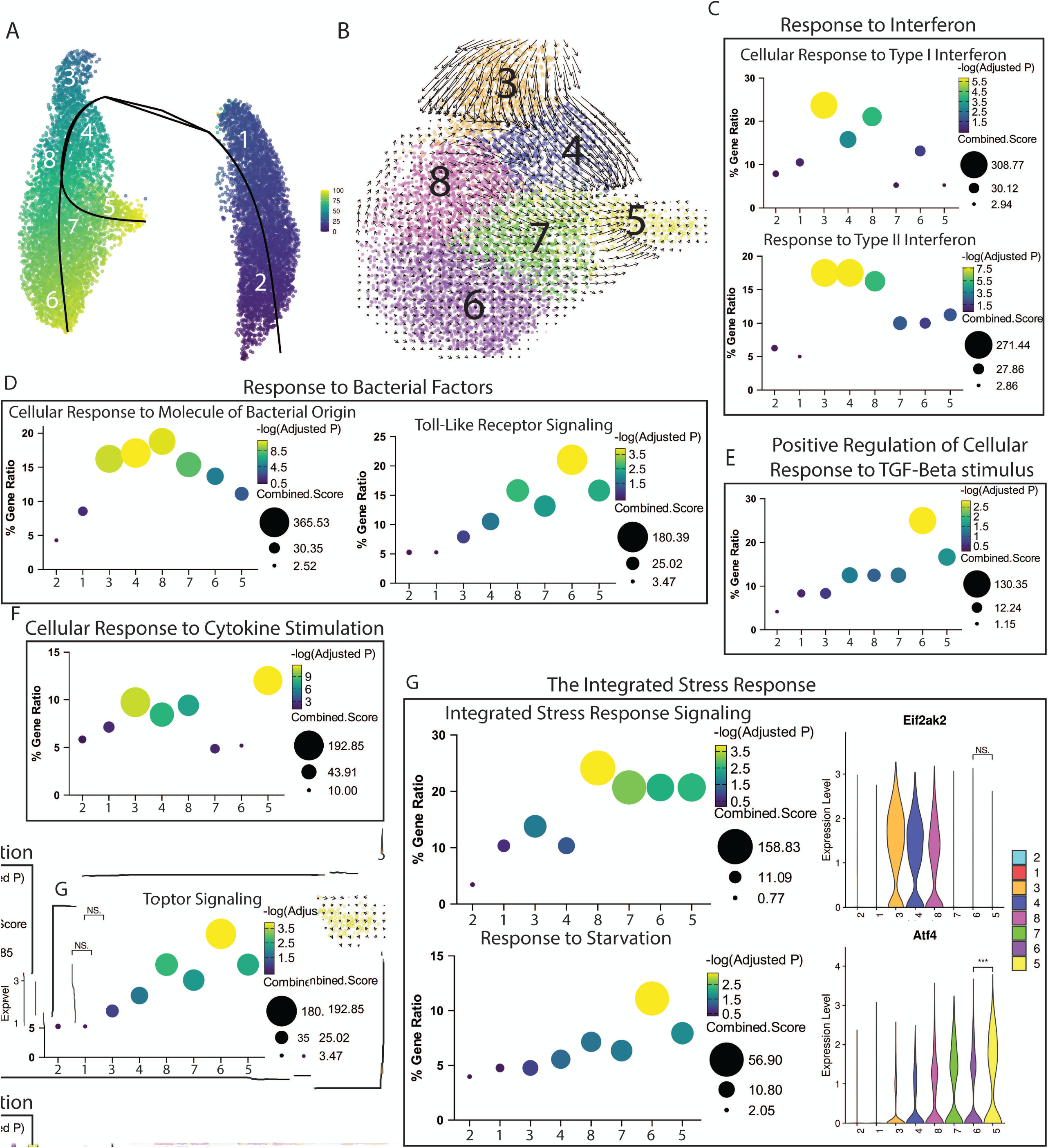
BALNs bifurcate into early and late phenotypes driven by discrete signaling responses. Unsupervised analyses were used to make inferences about the trajectory of RNA expression by neutrophils during pneumonia. (A) Slingshot was used to show pseudotime and bifurcation of RNA transcription trajectories from eight blood and BAL neutrophil clusters. (B) RNA velocity analysis performed using only BALNs shows localized changes in RNA expression. (C-G) Specific signaling pathways with neutrophil clusters (Slingshot pseudotime order) versus % gene ratio, –log_10_(Adjusted P), and combined score (-log(p)*odds ratio). Pathways: (C) response to interferon, (D) bacterial factors, (E) TGF-β, or (F) cytokine stimulation. (G) Plot of the integrated stress response (ISR) and response to starvation, with *Eif2ak2* and *Atf4* violin plots by cluster. For violin plot statistics comparing clusters 5 and 6, *** = FDR *q* < 0.001, NS = no significance.

DEGs from each cluster were used to perform gene ontology (GO) enrichment analysis to determine which pathways are overrepresented relative to each neutrophil cluster. The top ten pathways enriched from each cluster are presented in a bubble plot showing the gene ratio and adjusted P-value (**Figure 2E**). Blood C1 is enriched in translation machinery transcripts while blood C2 is enriched for pathways involved in granule content production and the electron transport chain, further suggesting that blood neutrophil transcription occurs in two distinct phases (**Figure 2E**). Of note, some functional pathways such as ficolin-1-rich and secretory granule production are active in both the BAL and blood clusters, suggesting that some BALN subsets may require additional granule production once in the infected airspace (**Figure 2E**). BALN C5 also has peptide biosynthesis, ribosome, and translation pathways enriched, suggesting that this group may be uniquely oriented for protein production in the airspace (**Figure 2E**). Interestingly, blood clusters are not significantly enriched for pathways relating to response to pathogens including molecules of bacterial origin, lipopolysaccharide, or inflammatory response. This suggests that blood neutrophil transcription is organized to facilitate maturation and migration and is not oriented for enabling neutrophil effector function. Taken together, these data suggests that blood neutrophil transcription serves drastically different purposes than BALN transcription in the infected airspace.

### BALNs bifurcate into early and late phenotypes driven by discrete signaling responses

To define how the neutrophil cell clusters relate to each other, we performed trajectory and RNA velocity analyses (28, 29). The trajectory identified by the Slingshot algorithm starts in the blood C2, passes through blood C1, then BAL C3, and bifurcates to mature to either C5 or C6 with C4, C7 and C8 as intermediate or transient populations (**Figure 3A**). RNA Velocity of BALN only (**Figure 3B**) or all neutrophils (**Supplemental Figure 2C**) also shows a strong flux of RNA expression running from C3 to C5, and to a more subtle extent from C3 to C8 and then from C8 to C6. Based on this trajectory data and gene fingerprints (discussed below), we will refer to cells in C3 as “early BALN” neutrophils and C5 and 6 neutrophils as “late BALN.” We next explored the pathways that differentiate these important clusters in more detail. GO enrichment from all DEGs from each cluster are shown with the top 20 pathways specifically upregulated in late BALN C3, C5, and C6 (**Supplemental Figure 2D-F**) and the full list of enriched pathways for each cluster is also presented (**Supplemental Table 4)**. Specific pathways of interest relevant for neutrophil function are presented and analyzed further in the next section.

To determine which cell phenotypes or gene pathways underlie the various BALN clusters, we manually selected and present signaling pathways key for neutrophil function that were enriched in BALN clusters via GO enrichment. A selection of these pathways is depicted in **Figure 3C-G** with clusters organized in the Slingshot-associated order. A full list of genes contributing to all enriched GO pathways is presented in **Supplemental Table 4**. Several pathways appear to distinguish the early from the late BALN clusters. Early and intermediate BALN clusters C3, C4, and C8 are highly enriched for responses to both type I and II interferon while late BALN C5, C6 and C7 have a low to intermediate response to type I and II interferon (**Figure 3C**). Early BALN cluster C3 is also highly enriched for cellular response to molecule of bacterial origin while late BALN cluster C6 is the highest enriched cluster for Toll-Like receptor signaling (**Figure 3D**). Curiously, C6 is the only cluster enriched for positive regulation of cellular response to TGF-β stimulus and C5 but not C6 is highly enriched for response to cytokine stimulation, suggesting that these signaling pathways may drive differences in transcriptional phenotypes (**Figure 3E-F**). Lastly, the integrated stress response (ISR) appears to be turned on in early BALNs and increases in late BALNs (**Figure 3G**). The ISR is activated via phosphorylation of eIF2α through several kinases including PKR (*Eif2ak2*), which results in the inhibition of cap-dependent mRNA and ATF4-dependant gene expression (30). Interestingly, *Eif2ak2* is expressed in early BALN clusters while *Atf4* and downstream genes are expressed in late BALNs. The response to starvation which is also *Eif2ak2-* and *Atf4*-dependent (31), is highly enriched in C6 only in our data (**Figure 3G**). Taken together, these data suggest that early BALNs are enriched in transcripts associated with response to viral and bacterial factors, while late BALNs bifurcate into either a C5 phenotype that is predominantly cytokine response driven, or a C6 phenotype that is influenced by TLR and TGF-β signaling. Though late BALNs are universally decreased in many pathogen-responsive pathways, they have dramatically disparate gene expression patterns, which will be explored more below.

### Late BALN subsets represent neutrophils with different effector functions

We next explored pathways that differentiate late BALN C5 and C6, starting with pathways specific to neutrophil effector functions. The production of granules, degranulation and phagocytosis are central processes in neutrophil pathogen clearance. The transcription of components of Azurophil and Ficolin-1-rich granules is strong in C6 (**Figure 4A**), while Specific, Tertiary, and Secretory granule components are highly transcribed in blood neutrophils but not BALNs (**Supplemental Figure 3A**). It has previously been reported that generation of neutrophil granules occurs exclusively in bone marrow and blood neutrophils, and not during or after migration to infected tissue (32, 33). The upregulation of granule-associated genes in C6 may indicate a second wave of granule production in mature neutrophils. C6 BALNs are also highly enriched for phagocytic machinery, including phagocytic vesicles, lysosome, and vacuolar proton-transporting V-type ATPase complex (**Figure 4B**), suggesting that phagocytosis is a key role for these cells.

**Figure 4.**
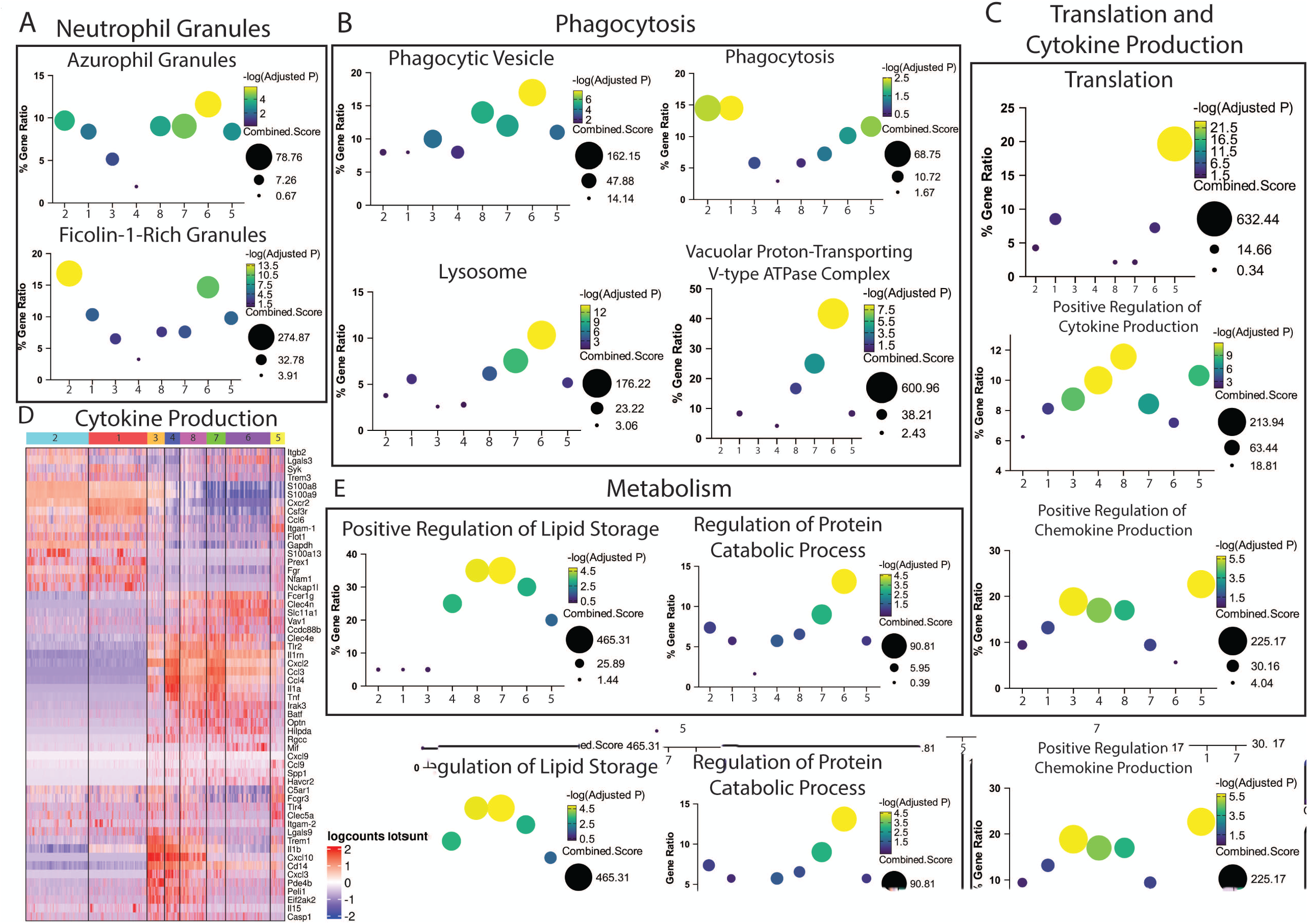
Late BALN subsets represent neutrophils with different effector functions. Specific canonical neutrophil function pathways plotted by neutrophil clusters (Slingshot pseudotime order) versus % gene ratio, –log_10_(Adjusted P), and combined score (-log(p)*odds ratio). (A) neutrophil granules, (B) phagocytosis, (C) translation and cytokine production, and (E) metabolism. (D) Heatmap of cytokine and associated gene expression for each cluster. Rows and columns correspond to genes and cells, respectively; columns are subdivided into Celda clusters; colors are scaled by row, with blue and red indicating values that are at least 2 standard deviations below or above the mean (white), respectively.

Another important function of neutrophils is the production of inflammatory cytokines and chemokines, which appears to be highly active in C5 but not C6 (**Figure 4C**). Pathways involved in ribosomal machinery, translation, and cytoplasmic translation are activated in C5, suggesting this machinery may be needed for prolonged cytokine production (**Figure 4C; Supplemental Figure 3B**). Additionally, distinct transcription factors that regulate inflammatory cytokine production are expressed significantly higher in C5 compared to C6, including *Irf5, Nfat5, Stat1, Stat3, Jak2*, as well as type 1 interferon receptor genes *Ifngr1* and *Ifngr2* (**Supplemental Figure 3C-D**). STAT1 and STAT2 are necessary for the response to interferon, while STAT3 and JAK2 are both known to mediate IL-3, IL-6, and IL-10 family cytokine responses (34, 35). C5 also transcribes high levels of the associated cytokine receptors *Il6ra, Il10ra, Il17ra, Il4ra,* and *Csf3r* (**Supplemental Figure 3E**). Interestingly, both late BALN clusters express chemokines *Cxcl2, Ccl3,* and *Ccl4*, though early and intermediate BALN clusters express these to a greater extent while also expressing *Cxcl10* and *Cxcl3* (**Figure 4D**). In terms of regulation, the role of IL-1 signaling appears to affect C6 to a greater extent than C5. IL1 signaling has been shown to be a key activator of neutrophils that can induce degranulation and NETosis (36, 37). C3 and C5 but not C6 express *Il1b*, while all BALNs express the receptor antagonist *Il1rn* (**Figure 4D**).

Curiously, C3 and C5 are the only BALNs that express the decoy receptor *Il1r2* (**Supplemental Figure 3E**). This may suggest that C6 is susceptible to IL-1 signaling but early and late C5 are resistant due to antagonistic and decoy IL-1R gene expression. Overall, this data regarding neutrophil effector function suggests that C5 BALN are driven by inflammatory cytokine signaling possibly via IL-6, IL-10, IL-17, IL-4, or G-CSF and function as professional cytokine producers while C6 BALN are driven by response to bacterial factors, TGF-β, and potential IL-1 signaling, leading to upregulating production of phagocytic machinery.

### Late BALN subsets have altered regulation of metabolic pathways and may have divergent cell fates

Upregulation and reliance on glycolysis for energy production has long been considered a hallmark of mature neutrophils, though evidence from non-lung infection models in the last decade also suggests that neutrophils can also utilize the TCA cycle, fatty acid oxidation (FAO), oxidative phosphorylation (OXPHOS), and the pentose phosphate pathway (PPP) to facilitate effector functions (38–42). Curiously, C6 but not C5 is enriched for positive regulation of lipid storage, and regulation of protein catabolic processes (**Figure 4E**), though FAO and PPP are not enriched in our dataset (**Supplemental Table 4**). To assist in determining which clusters are most strongly associated with specific pathways and gene sets, Variance-adjusted Mahalanobis (VAM) scoring was performed using curated gene sets adapted from Xie et al. 2020 and Cowland and Borregaard 2016 (**Supplemental Table 5**) (43–45). Reactive oxygen species (ROS) can be generated either via glycolysis or the mitochondrial electron transport chain. C6, but not C5 is highly enriched for glycolysis, which is driven by *Pkm, Eno1, Aldoa,* and *Pgam1* expression (**Supplemental Figure 4A**), in agreement with our prior data that C6 is enriched for phagocytosis associated pathways, and the current literature which reports that phagocytosis is dependent on glycolysis (38). Both late BALN clusters have comparably high VAM scores for the electron transport chain, though C6 is the highest enriched cluster for ROS production which appears to be driven by *Cyba, Sod2,* and *Ncf1* expression (**Supplemental Figure 4B-C**).

Lastly, to determine whether neutrophil cell fates are distinct among the BALN clusters, we investigated differences in cell death pathways within our scRNAseq dataset. All BALN clusters were enriched for positive regulation of programmed cell death (**Supplemental Figure 4E**). A crucial bactericidal cell death pathway for neutrophils is NETosis. Assessing NETosis via transcriptomics data has proven challenging, however recent efforts by Li, X., et al. 2023 have produced pro-NETosis associated gene sets that can be used to interpret whether neutrophils are likely to NETose (46). C5 has the highest NETosis VAM score, while C6 has the lowest (**Supplemental Figure 4D**). Cluster C6 is also the group most enriched for response to TGF-β (**Figure 3E**), and C6 but not C5 is active for positive regulation of Angiogenesis (**Supplemental Figure 3F**), as well as genes that quench digestive enzymes released from degranulation such as *Slpi* (**Supplemental Figure 3G**). Taken together, this suggests that although C6 BALNs are pro-phagocytic, they may also have a compensatory pro-resolution and tissue repair function, whereas C5 neutrophils secrete inflammatory cytokines and may promote NETosis.

### Neutrophil transcriptome is distinct between the pulmonary vasculature, interstitial, and alveolar space compartments

To understand how migration and location might affect neutrophil transcription, we next investigated changes that occur in the neutrophil transcriptome as they transit from the circulation to the alveolar space. Microarray analysis was performed on pneumonic neutrophils that were FACS-sorted from the lung vasculature, interstitium, and alveolar space (**Figure 5A**). BALF neutrophils were also collected and analyzed but were found to have very similar phenotypes to alveolar neutrophils that remained in the lung after lavage and were excluded from additional analysis (**Supplemental Figure 5A**). Overall, there were 7755 DEG across the three groups (one-way ANOVA after expression filtering, FDR *q* < 0.05), which were stratified into six clusters by hierarchical clustering (**Figure 5B. Supplemental Table 6)**. Compartment cluster 1 (CC1) is comprised of a small group of genes that are expressed most highly in the interstitial compartment (**Figure 5B and 5E**). The remaining clusters can then be described and organized into metagroups based on the pattern of their regulation. Metagroup 1 (MG1) is comprised only of CC1. Clusters with genes increased in the alveolar space (metagroup 2, MG2) include compartment cluster 6 (CC6, “late-up”), with genes that are only upregulated in alveolar neutrophils; compartment cluster 4 (CC4, “gradual-up”), upregulated in the interstitium and further upregulated in the alveolar airspace; and compartment cluster 5 (CC5, “early-up”), upregulated in the interstitium and remains highly expressed in the alveolar airspace (**Figure 5B and 5D**). Similarly, metagroup 3 (MG3) has clusters with genes decreased in the alveolar space and is comprised of compartment cluster 3 (CC3), described as “early-down” and compartment cluster 2 (CC2), described as “late-down” (**Figure 5B and 5E**).

**Figure 5.**
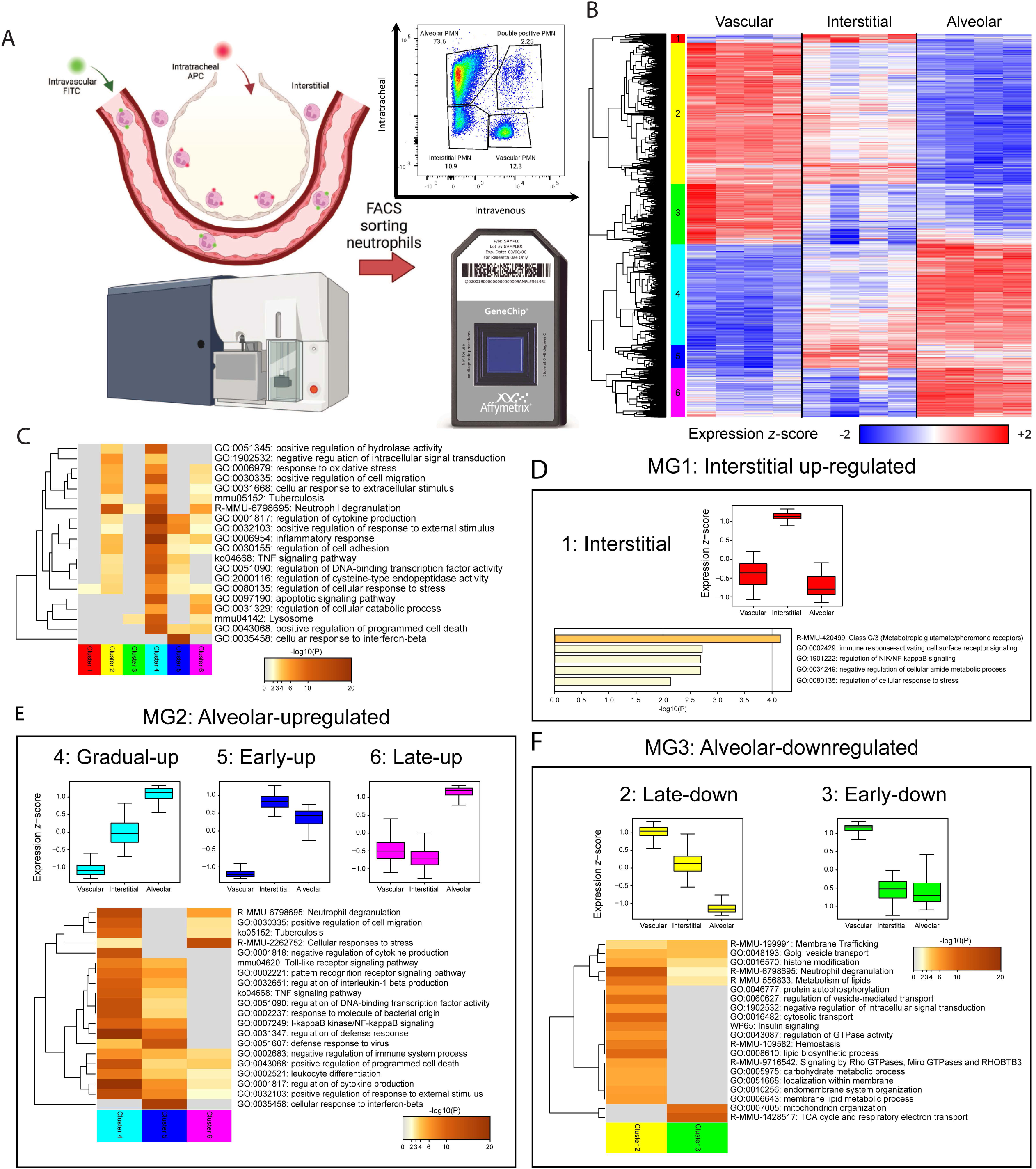
Neutrophil transcriptome is distinct between the pulmonary vasculature, interstitial, and alveolar space compartments. Four female mice were infected with ∼ 1.0×10^6^ CFU *S. pneumoniae* or saline control for 24h. (A) Experimental schematic. FITC-CD45 was instilled retro-orbitally to stain intravascular neutrophils prior to sacrificing the mouse, then intratracheal neutrophils were stained via APC-CD45 antibody. Neutrophils were isolated via FACS and profiled by Affymetrix gene expression microarray. (B) Heat map of DEGs (one-way ANOVA FDR *q* < 0.25, |fold change| > 2 between any two groups, n=3114 genes), divided into six clusters by hierarchical clustering. Rows and columns correspond to genes and samples, respectively; colors are scaled by row, with blue and red indicating values that are at least 2 standard deviations below or above the mean (white), respectively. (C) Metascape was used to compute pathway enrichment within each cluster; top enriched pathways were grouped hierarchically in a heatmap of all six gene clusters. (D-F) Metascape analysis was re-run on metagroups (MG) to show pathways specifically enriched in (D) interstitial-upregulated, (E) alveolar-upregulated, and (F) alveolar-downregulated gene sets. Boxplots denote the relative expression of the genes in each cluster: expression values were *z*-normalized (to a mean of zero and standard deviation of one) across all 12 samples, and the mean *z*-score was computed for each gene across the samples in each group. The boxes indicate the median and first and third quartiles, and the whiskers extend to the most extreme value within 1.5 times the interquartile range from the box.

To evaluate the differences between the gene clusters, we utilized Metascape (47) to determine which pathways are overrepresented in each cluster (**Figure 5C and Supplemental Figure 5B-C**). CC2, CC4 and CC5 are enriched in genes associated with response to pathogens, inflammatory response, cytokine production, and regulation of migration. These clusters are dynamically changed around the interstitial compartment, suggesting that transcriptional remodeling for numerous pathways occurs or begins in the interstitial space (**Figure 5C**). Next, an enrichment network was generated to show interactions between upregulated pathways where the circles represent pathways, sized in proportion to the number of genes, and subdivided by the proportion of each cluster comprising the node (48). Interestingly, the cellular response to stress is enriched in all clusters except for CC3 (**Figure 5C**), and is centrally located in the enrichment network, connecting numerous pathways including negative regulation of intracellular signaling, cytokine production, apoptotic signaling, and response to oxidative stress (**Supplemental Figure 5B-C**). These data may indicate that key neutrophil transcriptional regulation occurs in the pulmonary interstitium prior to migration to the airspace where the integrated stress response (ISR) may be a critical process.

### Neutrophil metabolism and ISR transcriptome are rewired prior to migration to the airspace

To further identify pathways associated with the various compartments, we used Metascape to analyze pathways specific to our metagroups. MG1 (CC1) is associated with a small number of pathways that are upregulated in the interstitium, including pheromone receptors and regulation of cellular response to stress (**Figure 5F**). MG1 genes include *Eif2ak2, Gabbr1*, and a group of G-coupled protein receptors, taste receptor type 2s (*Tas2r126, Tas2r135* and *Tas2r143*). *Gabbr1* and the TAS2Rs have been implicated as potential regulators of neutrophil function (49–51).

Within MG2 (CC4, CC5, CC6), many shared pathways pertain to antimicrobial response, especially between CC4 and CC5 (**Figure 5D**). Cellular response to IFN-β is almost exclusively associated with CC5, which may limit tissue damage as neutrophils migrate into the lung. Conversely, the cellular response to stress is most strongly associated with CC6, suggesting that something may be triggered in the interstitium that activates the ISR (**Figure 5D**). These data suggest that numerous pathways related to pathogen response, inflammation, and response to stress are transcribed as neutrophils migrate to the airspace, and that the interstitial space may represent a crucial intermediate point.

Pathways enriched in MG3 (CC2, CC3) include lipid and carbohydrate metabolism, TCA cycle, and respiratory electron transport chain (**Figure 5D**), which is consistent with the earlier results showing that neutrophils undergo metabolic rewiring as they migrate to sites of infection (**Figures 1 and 2**). Neutrophils are known to shut off respiratory pathways and rely on glycolysis for energy production in infected tissues, although recent work has shown that the TCA cycle can also be used to supply energy for NETosis (38). TCA cycle and mitochondrial genes are downregulated early, while lipid and carbohydrate metabolism and GTPase signaling are downregulated later (**Figure 5D**). This implicates a stepwise metabolic transition whereby neutrophils from the vasculature undergo downregulation of TCA and ETC as they migrate to the interstitium, then turn off lipid and carbohydrate metabolism upon migration to the alveolar airspace.

### EIF2AK2 expression is upregulated in the pulmonary interstitial compartment

Since a central theme of the DEGs between blood and alveolar neutrophils is regulation of cellular machinery, cells surrounding and within the interstitium are enriched for ISR pathways, and a regulator of the ISR, *Eif2ak2*, is specifically upregulated in neutrophils in the interstitium, we decided to investigate the ISR further in our model system. We first measured transcriptional changes of several CC1 genes by performing qRT-PCR of TAS2Rs*, Gabbr1,* and *Eif2ak2* in a parallel infection and sorting experiment. Curiously, only the expression of *Eif2ak2* is significantly higher in neutrophils from the interstitial compartment compared to those from the pulmonary vasculature or alveolar space (**Figure 6A; Supplemental Figure 5D**). To provide spatial context to neutrophil gene expression in the pneumonic lung, RNAscope was performed on mice that had been infected for 24-hours with SP3. *Eif2ak2* puncta (red) were identified in *S100a8*+ (green) neutrophils, which were classified as residing in large blood vessels, peri-vascular interstitium, alveolar septum, or alveolar airspace (**Figure 6B**). *Eif2ak2*+ neutrophils with two or more red puncta represented a significantly higher fraction of neutrophils in the peri-vascular interstitium compared to those in all other compartments (**Figure 6C**). We also investigated *Eif2ak2* and the downstream transcription factor *Atf4* in our scRNAseq dataset. Consistent with these results, early BALN (C3) express higher levels of *Eif2ak2* compared to blood (C1, C2) or late BALN (C5, C6), and *Atf4* expression is higher in intermediate and late BALNs (C5-C8) (**Figure 3G**). Further, in our scRNAseq GO enrichment, the ISR is significantly enriched in early and late BALN but not blood clusters (**Figure 3G**). Taken together, these results indicate that *Eif2ak2* is distinctly transcribed in the pulmonary interstitium and could represent a previously unreported crucial step in neutrophil transcriptional regulation before migration to the alveolar airspace (**Figure 6D**).

**Figure 6.**
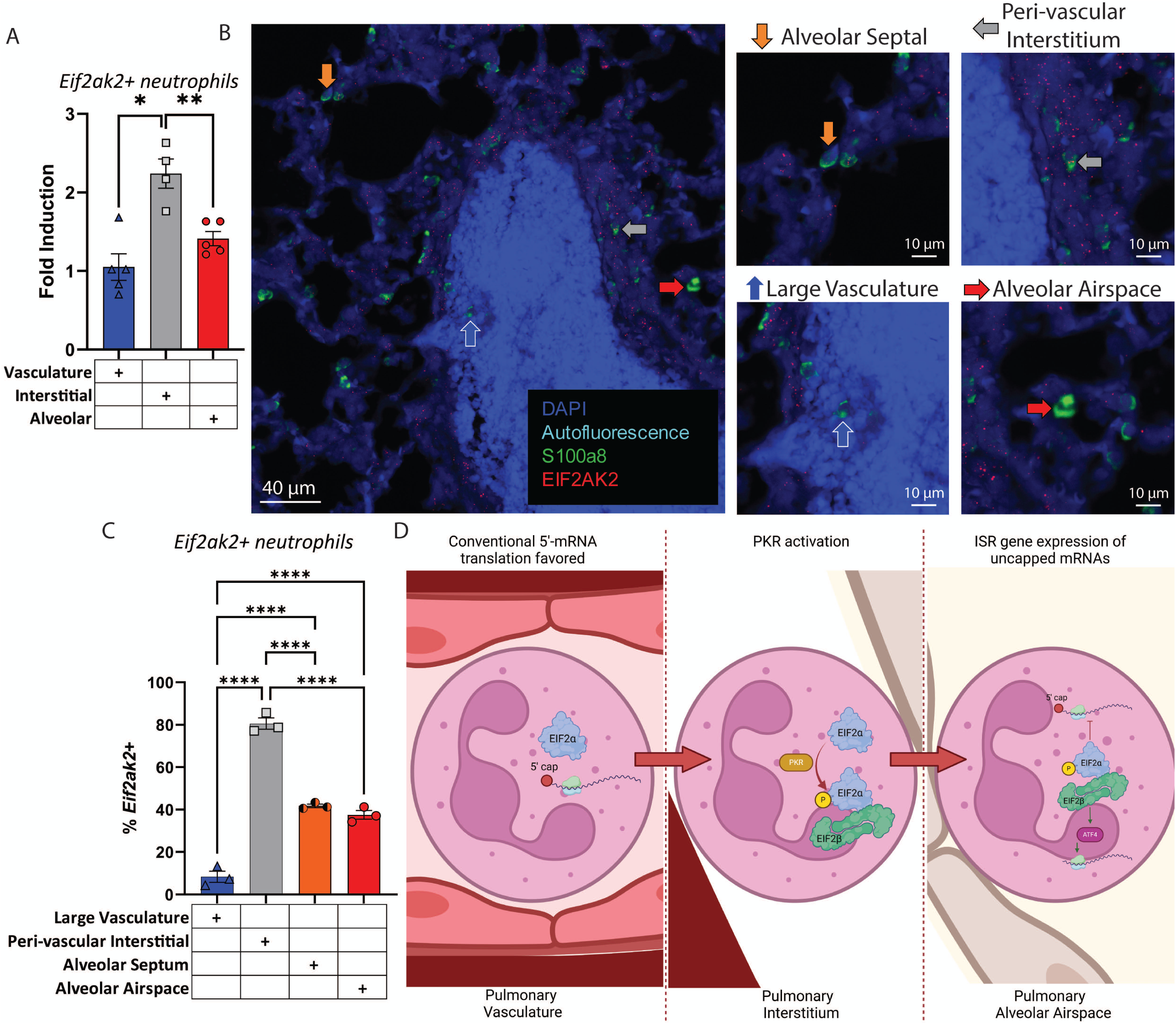
EIF2AK2 expression is upregulated in the pulmonary interstitial compartment. (A) Neutrophils from SP3-infected mice isolated from the pulmonary vasculature, interstitium, or airspace sorted by FACS and qRT-PCR was performed on *Eif2ak2* and normalized to expression from neutrophils from the pulmonary vasculature from uninfected mice. (B) Paraformaldehyde fixed paraffin embedded SP3-infected lungs were cut into 10µm sections and RNA-ISH (RNAscope) was performed for *S100a8* (neutrophils, green) and *Eif2ak2* (red), with nuclei labeled with DAPI (blue) and red blood cells identified via autofluorescence. Representative images shown, with arrows denoting neutrophils in alveolar septum (orange), peri-vascular interstitium (gray), large vasculature (blue) and alveolar airspace (red). (C) *Eif2ak2*+ (≥ 2 puncta) were quantified by pulmonary compartments which included the pulmonary large vasculature, peri-vascular interstitium, alveolar septum, and alveolar airspace. (D) Schematic representation of hypothesized role of PKR in the ISR rewiring transcription in neutrophils as they migrate from the peripheral blood and pulmonary vasculature through the interstitial space to the alveolar airspace. Statistics presented are one-way ANOVAs with Bonferroni multiple comparison corrections where the adjusted P value is represented by * = P < 0.05, ** = P < 0.01, and **** = P < 0.0001.

## Discussion

Here we describe various aspects of the transcriptome of neutrophils as they travel from the blood to the airspace during pneumococcal pneumonia. While other studies have investigated the transcriptomics of lung neutrophils in other infection models, ours is the first such study that we are aware of that focuses on lung neutrophils during bacterial pneumonia. During infection, the transcriptome of circulating neutrophils differed dramatically from that of alveolar neutrophils yet remained similar to that of circulating neutrophils in uninfected mice (**Figure 1B-C**).

Alveolar neutrophils were characterized by strong upregulation of genes associated with cytokines and inflammation and downregulation of genes associated with cellular machinery, and could be divided into distinct early and late BALN phenotypes: early BALNs that respond to interferons and bacterial factors (**Figure 3A-G**), and two subsets of late BALNs that either respond to type I interferon to produce cytokines and chemokines and are pro-NETosis or respond to IL1 signaling and bacterial factors and activate degranulation and phagocytosis pathways (**Figure 3F-I and 4A-E**). Evidence of neutrophil metabolic reprogramming is also shown, as neutrophils begin to shut off TCA, ETC, and lipid metabolism in the pulmonary interstitial space (**Figure 5E**), but specific subsets of late BALNs upregulate ETC (C5 and C6) (**Supplemental Figure 3C**) and glycolysis (C6) (**Figure 3A**). The integrated stress response (ISR) is implicated as a key regulatory “switch” that may facilitate neutrophil transcriptional rewiring as neutrophils migrate to the pneumonic lung. EIF2 signaling is downregulated and PKR in IFN induction is upregulated in BALN compared to blood neutrophils (**Figure 1E**), the ISR is enriched in late BALNs (**Figure 3J**), and Eif2ak2 is highly expressed in interstitial neutrophils (**Figure 5B-D**).

In recent years, murine and human studies have shown that neutrophil transcriptome and function vary based on the tissue of origin (12, 52). Our data agree with the literature, and additionally show that neutrophil location affects the transcriptome more than the overall infection status of the mouse (**Figure 1A-D**). The exact environmental cues that drive the changes in neutrophil transcriptome are not fully clear. At homeostasis, Ballesteros et al. suggest that CXCL12-CXCR4 interactions are required for a pulmonary-specific lung neutrophil transcriptome. In our model, we see that late BALN C5 (but not other neutrophil clusters) highly express *Cxcr4* (**Figure 2D**), which may contribute to NF-κB driven transcription, though numerous other cytokine-mediated signaling pathways may do so as well. At homeostasis, whole-lung neutrophils exhibit angiogenic characteristics that neutrophils from other tissues lack (12). Interestingly, our work also shows that a specific subset of late-degranulating BALNs express genes involved in positive regulation of angiogenesis (**Supplemental Figure 3F**). Future proteomics work on later timepoints in sublethal pneumonia models will be needed to elucidate the role of neutrophils and angiogenesis post-acute lung injury, and to validate the functional status of these pneumonic neutrophils.

We demonstrate multiple distinct populations of alveolar neutrophils during pneumonia, which we have characterized as early BALN (C3), late-degranulating BALN (C6) and late-cytokine producing BALN (C5). These subsets are consistent with aspects of prior work and add additional information to our knowledge of neutrophils. Recent single cell studies of human and murine peripheral and bone marrow neutrophils demonstrate two groups of mature neutrophils that stratify by IFN-inducible and non-IFN-induced gene expression profiles (44, 52, 53). Similarly, in our scRNAseq data, C3 is enriched for response to type I and type II IFN, and C6 (but not C5) is enriched for response to type I interferon (**Figure 3C)**. Our cytokine producing BALN C5 is also similar to neutrophils found in prior studies; specifically, immature (low density) neutrophils tend to produce inflammatory cytokines but have a limited capacity for respiratory burst and NETosis *in vitro* (54), and in *Cryptococcus neoformans* pulmonary infections, neutrophils were shown to adopt either a cytokine or an oxidative stress transcription signature (55). Contrary to these studies, however, our C5 BALNs may also be pro-NETosis (**Supplemental Figure 3D**).

It is not clear how the two discrete late BALN populations form. One possibility is that at 24 hours post infection, there is a mix of mature neutrophils marginated in the pulmonary vasculature prior to infection, and another group of neutrophils that were generated during emergency granulopoiesis and thus may be more primed for an inflammatory response. Recent transcriptional work in human sepsis patients has shown that during emergency granulopoieses, an immature neutrophil phenotype associated with STAT3 mediated production of IL-6 family cytokines and high expression of IL1R2 may drive disease (56). Taken together, this may suggest that our late BALN C5, which is also enriched for *Stat3, Il1r2,* and inflammatory cytokine production, is a phenotype resulting from emergency granulopoiesis, while C6 may be an aged phenotype. Future work should be performed to determine whether this C5 phenotype is a direct result of emergency granulopoiesis and is directly associated with host tissue damage that drives disease progression.

The impact of neutrophil metabolism on pneumonia disease progression is an understudied area. Our blood and BAL neutrophil microarray data show that fatty acid oxidation (FAO), oxidative phosphorylation (OXPHOS), and the TCA cycle are strongly downregulated as neutrophils migrate to the infected lung (**Figure 1G and 5F**). These data are consistent with and add to the growing volume of work that shows that neutrophils may use multiple sources of ATP besides glycolysis to facilitate effector function in infected tissues (38). Interestingly, our scRNAseq data also suggests that metabolic regulation is dynamic among alveolar neutrophils. Both late BALN clusters upregulate ETC (**Supplemental Figure 4C**), while pro-degranulation (C6) late BALNs upregulate glycolysis and C5 BALNs do not (**Supplemental Figure 4A**). This curious difference in the metabolic signatures between distinct mature neutrophil phenotypes may be a result of differences in effector function and cell-death-related mitochondrial function. C6 scores the lowest for NETosis while C5 scores the highest, suggesting that late BALN clusters have drastically different cell fates (**Supplemental Figure 4D**). Neutrophil NETosis can be induced in either a NADPH oxidase (NOX)-dependent pathway where glucose is required for cytosolic ROS production and leads to PAD4 activation, or a NOX-independent mechanism where mitochondria are necessary for ROS production and subsequent PAD4 activation (57). Further, murine BAL neutrophils after LPS challenge are known to scavenge extracellular proteins and use glutamine to produce TCA intermediates that can be used for energy production required for NETosis (41, 58). Therefore, late BALN C5 may facilitate NETosis through multiple metabolic avenues, some of which may not be clear from predominantly transcriptional analyses.

Interestingly, pulmonary compartment microarray data showed that interstitial neutrophils have decreased TCA and ETC (but not lipid and carbohydrate metabolism) compared to vascular neutrophils, while all these processes are downregulated in alveolar neutrophils (**Figure 5F**), which is in contrast to the dynamic regulation seen in our scRNASeq data. Likely, the signal from a relatively small number of pro-cell-death neutrophils was obscured by the other BAL neutrophils in a bulk analysis. Another possibility is that neutrophil metabolism is rewired as a result of neutrophil maturation; previous studies have shown that immature neutrophils retain the use of oxidative phosphorylation and FAO, while mature neutrophils do not use these pathways for energy production (59, 60). Our data comports with existing literature relating to non-lung tissue and *in vitro* data, since immature (but not mature) neutrophils are known to produce energy via lipophagy and directing fatty acids to the TCA cycle and oxidative phosphorylation (39, 59). These data suggests that neutrophils mature as they migrate to the pulmonary airspace in a highly organized step-wise fashion. Recent studies have shown that neutrophils can also use the pentose phosphate pathway (PPP), OXPHOS, and FAO to facilitate de granulation, NET production, and ROS production (38, 61–63). Taken together, these data indicate that neutrophil metabolism is tightly regulated as they migrate into the infected airspace to facilitate changes in maturation and effector function.

Integrated stress response (ISR) pathways are key to integrating environmental stresses with appropriate cellular responses. In one arm of this response, *Eif2ak2* (PKR) phosphorylates EIF2α, which binds to EIF2β, transitioning from cap-dependent to cap-independent translation, increasing production of the transcription factor ATF4, leading to increases in genes involved in interferon, anti-viral, and anti-bacterial responses (64–66). We believe that the PKR-induced ISR may play a central role in regulating the neutrophil transcriptomic patterns we see in our model of bacterial pneumonia. This is supported by data from all three of the presented transcriptomic studies. In blood versus alveolar neutrophils, canonical EIF2 signaling is the most significantly downregulated pathway and PKR in IFN induction and antiviral response is a significantly upregulated pathway (**Figure 1D**). This is consistent with previous studies showing that EIF2 signaling is downregulated in neutrophils during or at sites of infection (25, 67). Additionally, among BALN *Eif2ak2* is highest in early and intermediate BALN, while *Atf4* is highest in late BALN suggesting dynamic regulation of these pathways among neutrophils (**Figure 3G)**. Finally, to our knowledge, we are the first to report that *Eif2ak2* was increased specifically in interstitial neutrophils compared to vascular or alveolar neutrophils (**Figure 6A-C**). It is interesting that PKR-induction of the ISR is implicated in our *S. pneumoniae* model, as PKR was initially characterized by its response to dsRNA during viral infections. However, it has recently also been shown to respond to C5a (64, 68), which should be expressed at high levels in our model. Our data also suggests that the ISR and neutrophil metabolic status are intertwined. For example, PKR-ISR is upregulated in alveolar BAL neutrophils compared to peripheral blood neutrophils, with a corresponding metabolic reprogramming where oxidative phosphorylation, fatty acid oxidation, and the TCA cycle are downregulated (**Figure 1D**). Our findings agree with the literature in that ISR activation and ATF4 activation leads to the downregulation of TCA cycle enzyme translation (69). Recent work has also shown that during sepsis or LPS challenge, macrophages upregulate glycolysis via ATF4 activation which enhances effector functions such as inflammatory signaling and phagocytosis (70). This is consistent with our data that early BALNs express high *Eif2ak2* followed by late BALN (C6) *Atf4* expression and glycolysis transcription upregulation (**Figure 3G**). This link between ISR and metabolic reprogramming of neutrophils may thus be necessary to facilitate neutrophil maturation and subsequent effector function in the lung.

Although this work significantly adds to the current literature and provides insights into neutrophil transcriptome remodeling during pneumonia, it is limited to RNA techniques. Future work utilizing proteomic and metabolomic techniques should be performed to identify the precise nature of the relationship between the neutrophil ISR pathway, metabolome, and effector function during pneumonia. While we accounted for multiple comparisons in our statistical methods and our microarray and scRNAseq analyses platforms are rigorous, the possibility of some false-positives should be considered, as well as the relatively low number of samples (n=3-4 per group) in most of the analyses. The general possibility of discrepancies between high quality mRNA, protein produced, and actual neutrophil function should also be considered. We report discrepancies in some specific genes between microarray and qPCR generated data (**Supplemental Figure 5D**), though these are likely due to some genes within the 7755 DEG dataset appearing to have a sporadic signal in the interstitium. Our confidence in our conclusions is bolstered by the recurrence of transcriptomic patters across all three of the studies presented. Signaling pathway analysis is also difficult to interpret, since many signaling pathways are phosphorylation dependent, including the JAK/STAT, MAPK, and NF-κB pathways. For example, the absence of transcription factors like *Stat2* from specific clusters may indicate that this cluster does not have much if any STAT2 available to facilitate signaling, but the presence of STAT2 mRNA does not mean that that cluster necessarily relies on STAT2 activation. Proteomic, metabolomic, and functional studies should be performed in subsequent work. Lastly, we assume in this work that neutrophils in the interstitial compartment are transitory and en route to the airspace. To our knowledge, including our own unpublished data, resident interstitial neutrophils have not been reported during infection or during homeostasis. However, additional work will need to be done to further characterize the locational and temporal nature of these cells.

Accumulating evidence suggests that neutrophils are highly transcriptionally plastic and reprogrammed as they migrate to the pulmonary airspace during pneumonia. We demonstrate that the neutrophil transcriptome is drastically different between infected blood and BAL neutrophils and between neutrophils in various pulmonary compartments and reveal insights into plasticity and heterogeneity within the airspace compartment. These results offer novel insights into the murine neutrophil transcriptome during pneumococcal pneumonia, including how the ISR and metabolic pathways may play a critical role in neutrophil maturation and function. *Eif2ak2* stands out in our work as a likely regulatory switch during migration to the lung and a potentially desirable therapeutic target. Honing our understanding of neutrophil regulation though further proteomic and metabolomic work will be necessary to determine the importance and viability of targeting various neutrophil subsets during pneumonia.

## Methods

### Sex as a biological variable

Transcriptomic studies were performed on both male and female mice. Due to constraints in the number of mice that could be included in each study, only either male or female mice were included in specific microarray or scRNA-seq experiments, and some discrepancies between experiments may be a result of sex differences. Follow up higher powered studies should be performed to investigate sex-based differences.

### Murine model of pneumonia

C57BL/6J mice were purchased from Jackson Laboratories (Jax.org). Mice were between 6-12 weeks of age, numbers of male and female mice are specified in figure legends. Mice were housed in a barrier facility and provided food and water *ad libitum.* For pneumonia models, approximately 1*10^6^ CFU *Streptococcus pneumonia* (*S. pneumoniae*, serotype 3, strain catalog number 6303; ATCC) was intratracheally injected into the left lobe of mice as previously reported (71).

### Isolation of neutrophil RNA from FACS-sorted peripheral blood and BALF

Peripheral blood was harvested from the IVC via a heparinized syringe, red blood cells were lysed osmotically (PharmLyse Buffer, BD Biosciences), and cells were resuspended in FACS buffer. For BAL cells, and the heart-lung block was removed and suspended on a 25G blunt needle. The lungs were serially lavaged with a total of 10ml ice cold sterile lavage buffer (Hanks’ balanced salt solution [HBSS; Life Technologies, Invitrogen], with 2.7 mM EDTA disodium salt solution, 20 mM HEPES, 100 U/ml penicillin-streptomycin [Pen-Strep] [Sigma-Aldrich]). BALF cells were collected by centrifugation at 300G for 5 minutes and resuspended in FACS buffer. Cells were stained with 7AAD (BioLegend), CD45-FITC (clone 30-F11, BioLegend) and Ly6G-APC (clone 1A8, BioLegend) to identify live (7AAD^−^), single cell, neutrophils (CD45^+^Ly6G^+^), which were then isolated using a BD FACS ARIA II SORP. Sorted cells were washed with PBS and re-suspended in RNAProtect (Qiagen) and frozen at –80°C. RNA was isolated from frozen neutrophils using the RNeasy micro kit (Qiagen), following the manufacturer’s instructions. RNA was eluted in 14µl RNase-free water and stored at –80°C.

### Isolation of neutrophil RNA from lung compartments

Prior to euthanasia, mice were retro-orbitally injected with CD45-FITC (2.5mg in 100µl, clone 30-F11, BioLegend). After 3 minutes, mice were euthanized, the heart-lung block was removed and suspended on a 25G blunt needle. CD45-APC (2µg in 1ml PBS, clone 30-F11, BioLegend) was instilled intratracheally and allowed to dwell in the lungs for 3 minutes. The lungs were then serially lavaged with an additional 9ml ice cold sterile lavage buffer. BALF cells were collected as above, resuspended in FACS buffer, and held on ice. Meanwhile, lungs were digested to single cell suspension using the gentleMACS lung dissociation kit and dissociator per the manufacturer’s instructions (Miltenyi Biotec). BALF cells or lung digest cells were stained with Ly6G-APC-Cy7 (clone 1A8, BioLegend), CD45.2 (clone 104, BioLegend) and Zombie Aqua viability dye (BioLegend). Neutrophils (CD45.2^+^/Ly6G^+^) from BALF, and lung digest vasculature (CD45-IV^+^ CD45-IT^−^), interstitium (CD45-IV^−^ CD45-IT^−^) and alveolar space (CD45-IV^−^ CD-45IT^+^) were isolated by FACS. RNA from sorted cells was isolated as described above.

### Microarray analysis

RNA was extracted from single cell suspensions as described above. RNA quality control was performed via an Agilent Bioanalyzer (Agilent, Santa Clara, CA). Samples with an RNA integrity number (RIN) ≥ 7.5 were used. The Affymetrix GeneChip Mouse Gene 2.0 ST array platform was used for our analyses (Affymetrix, Santa Clara, CA) (72). For each experiment, all samples were processed together to mitigate batch effects. CEL files were normalized to generate gene-level expression values via the Robust Multiarray Average (RMA) in the affy package (version 1.58.1) and Entrez Gene-specific R packages (version 23.0.0) from the Molecular and Behavioral Neuroscience Institute (Brainarray) at the University of Michigan (73). Gene biotypes were obtained using the biomaRt R package (version 2.38.0). Array quality was assessed by computing Relative Log Expression (RLE) and Normalized Unscaled Standard Error (NUSE) using the affyPLM R package (version 1.58.0). Differential expression was assessed using the moderated (empirical Bayesian) ANOVA and *t* test implemented in the limma R package (version 3.38.3) Correction for multiple hypothesis testing was accomplished using the Benjamini-Hochberg false discovery rate (FDR) (74). Human homologs of mouse genes were identified using HomoloGene (version 68). All microarray analyses were performed using the R environment for statistical computing (version 3.5.1).

### Ingenuity Pathway Analysis (IPA)

Canonical pathways and upstream regulators were predicted from DEG via Ingenuity Pathway Analysis (IPA; version 01-16; Qiagen, Hilden, Germany) (19). A dataset composed of gene identifiers, fold change, and FDR-corrected *p* values (*q* values) was uploaded, and a *q* value threshold of 0.1 was set. Upstream regulators were further refined to a curated set of known transcription factors. Significance was calculated using a right/tailed Fisher’s exact test P value calculation. Z-scores were calculated using the consistency of pattern match of up– and down-regulated genes in the dataset and the expected activation and inhibition pattern downstream of a given regulator for both canonical pathway and upstream regulator analyses.

### Gene Set Enrichment Analysis (GSEA)

Entrez Gene IDs of the human homologs of all genes on the Mouse Gene 2.0 ST Array were ranked by the t statistic computed for each comparison. Any mouse genes with multiple human homologs (or vice versa) were removed prior to ranking, so that the ranked list represents only those human genes that match exactly one mouse gene. This ranked list was compared to the publicly available Molecular Signatures Database, or MSigDB (version 6.0). A p value was assigned to each gene set based on how skewed the distribution of the gene set is towards the up-or down-regulated end of the ranked list.

### qRT-PCR

RNA was isolated from cells as described above. RNA concentrations were determined using a NanoDrop spectrophotometer (Thermo Scientific, Waltham, MA). Specific RNA transcripts were quantified using a TaqMan RNA-to-CT 1-step kit and the QuantStudio 3 real-time PCR system (Thermo Scientific). All threshold cycle (CT) values were normalized to 18S rRNA. Expression values are presented as fold inductions after normalization. Primers and probes for specific gene targets are listed in **Supplemental Table 7** in the supplemental material.

### Single cell RNA sequencing (scRNAseq)

Peripheral blood and BAL cells were collected as described above, and resuspended in EasySep Buffer (Stemcell Technologies). Neutrophils were enriched using a negative selection magnetic bead enrichment kit (Stemcell Technologies). Sample viability and counts were determined using 10x recommended Countess II Automated Cell Counter. Single cells, reagents, and a single Gel Bead containing barcoded oligonucleotides are encapsulated into nanoliter-sized GEMs (Gel Bead-in-Emulsion) using the GemCode platform. Lysis and barcoded reverse transcription of RNAs from single cells is performed. Full length, barcoded cDNA was amplified by PCR to generated sufficient mass for library construction. Results can be viewed in the cDNA Bio Analyzer QC summary files. Enzyme fragmentation, A tailing, adaptor ligation and PCR are performed to obtain final libraries containing P5 and P7 primers used in Illumina bridge amplification. Size distribution and molarity of resulting cDNA libraries were assessed via Bioanalyzer High Sensitivity DNA Assay (Agilent Technologies, USA). All cDNA libraries were sequenced on an Illumina NextSeq 500 instrument according to Illumina and 10x Genomics guidelines with 1.4-1.8pM input and 1% PhiX control library spike-in (Illumina, USA).

### Preprocessing and quality control of single-cell data

The 10X CellRanger tool v3.1.0 was used for demultiplexing, alignment, identification of cells, and counting of unique molecular indices (UMIs). Specifically, the CellRanger mkfastq command was used to demultiplex raw base call (BCL) files generated by Illumina sequencers into FASTQ files. The CellRanger count command was used to perform alignment and create UMI count matrices using parameters –expect-cells=5000. Droplets with at least 500 UMIs underwent further quality control with the SCTK-QC pipeline (75). The median number of UMIs was 3,736 and 4,470 in BAL and blood samples, respectively, the median number of genes detected was 997 and 1,291, the median percentage of mitochondrial reads was 0.3% and 0.6%, and the median contamination estimated by decontX (76) was 0.25 and 0.18. Cells with less than 500 counts, less than 500 genes detected, or more than 5% mitochondrial counts were excluded. Cells were further filtered by visualizing canonical neutrophil marker genes, removing cells that appeared to be non-neutrophils, and leaving 11,037 cells for downstream analysis.

### Clustering of single-cell data with Celda

The celda package was used to bi-clustering genes into modules and cells into subpopulations (22). The top 5,000 variable features are selected with the runSeuratFindHVG function from the singleCellTK package, removing features with less than 3 counts in 3 cells. The recursiveSplitModule and recursiveSplitCell functions were used to select the model with 100 modules and 8 cell subpopulations after examining the Rate of Perplexity Change (RPC). Cells were embedded in two dimensions with UMAP using the celdaUmap function. Heatmaps for specific modules were generated using the moduleHeatmap function. Markers for each cluster were identified with the findMarkerDiffExp function from the singleCellTK package using a Wilcox rank sum test and an FDR threshold of 0.05.

### Pseudotime inference with Slingshot

The Slingshot package was used to infer pseudotime and cell trajectories (29). The slingshot function was applied using the celda cluster labels, without specifying a start or end cluster. The trajectory was embedded to the previously generated UMAP with the embedCurves function, then the slingCurves function was used to extract the trajectory curves for visualization.

### Estimation of RNA velocity

Loom files were created with loompy fromfq. The mouse index was built with the instructions provided on the loompy Github page. The velocity was estimated with the RunVelocity function with parameters kCells = 25 and fit.quantile = 0.02 from the Velocyto.R package (28). The velocity embeddings were then shown on the previously generated UMAP embedding.

### Enrichment Analysis of scRNA-seq Clusters

Gene set enrichment scores were calculated with the run VAM function from the singleCellTK package, using the log-normalized counts as input (43). Gene ontology term enrichment was calculated with the runEnrichR function from the singleCellTK package, using differentially expressed genes with log2 fold change > 0.25 in each cluster (77).

### Multiplex fluorescence RNA in situ hybridization (RNA-ISH)

Multiplex fluorescence RNA-ISH was performed using the RNAscope Multiplex Fluorescent Reagent Kit v2 (Advanced Cell Diagnostics, Inc., Newark, NJ, USA) according to the manufacturer’s instructions. Briefly, infected heart-lung blocks were inflated to 23 cmH2O and fixed in 10% neutral buffered formalin, and paraffin embedded. 10μm sections were deparaffinized, digested with protease, and hybridized for 2 hours at 40°C with the following RNAscope target RNA-specific probes: Mm-*Eif2ak2* (Cat. No. 822871) and MM-S100a8-C3 (Cat. No. 478511-C3). Preamplifier, amplifier, HRP-labeled oligos, and Opal 520 and Opal 570 dyes were then hybridized sequentially at 40°C. The nuclei were stained with DAPI, and images were acquired by Vectra Polaris PhenoImager HT 2.0 slide scanner and software (Akoya Biosciences). Two male and one female mice from two from two separate experiments were sectioned and quantified. *Eif2ak2* puncta were manually counted by two different operators. *S100a8*+ cells were identified, and the pulmonary compartment was classified before turning on the *Eif2ak2* channel. *Eif2ak2* status was then determined to be negative (0-1 puncta) or positive (2≤ puncta) for 500 (250 from each operator) random neutrophils across the entire infection-involved left lobe with roughly similar numbers from each compartment. *Eif2ak2*% positive neutrophils were compared by tissue compartment via one-way ANOVA with Bonferroni multiple comparison’s corrections. Operator’s counts had >95% agreement.

### Statistics

Unless otherwise stated, the statistical method used to compare multiple treatment groups was a one-way ANOVA with Bonferroni multiple comparisons corrections. Asterisks for adjusted p-values are * for p<0.05, ** for p<0.01, *** for p<0.001, and **** for p<0.0001.

### Study Approval

All animal studies presented in this manuscript were overseen by the Boston University Institutional Animal Care and Use Committee (IACUC). All protocols were approved prior to the performance of experiments (BU IACUC protocol #PROTO201800710).

### Data Availability

The raw and processed scRNAseq data has been deposited to GEO, with accession number GSE218681. Code used for analyses is available on Github https://github.com/scsdata/pihlr-traberk-neutrophils.

The microarray data discussed in this publication have been deposited in NCBI’s Gene Expression Omnibus and are accessible through GEO Series accession number GSE169007 (https://www.ncbi.nlm.nih.gov/geo/query/acc.cgi?acc=GSE169007).

## Author contributions

RMFP: experimental design, conducting experiments, data analysis, and manuscript writing and preparation, SHA: data analysis and writing, BEH: conducting experiments and writing, EMRA: conducting experiments, KRM: conducting experiments, ELD: conducting experiments, YL: data analysis and writing, JDC: experimental design, data analysis, and writing, ACG: data analysis and writing, JPM: experimental design and writing, LJQ: experimental design, writing and providing reagents, KET: experimental design, conduct experiments, data analysis, and writing.

## Acknowledgements

Funding sources:

KET: R01HL158732, K08130582, L30 HL138777, F32 HL120551, T32 HL703547, KL2

TR001411, UL1 TR001430 (BU-CTSI Sequencing Voucher)

JPM: NIH R01 HL171499, R01 AI162850, R01 AI115053, R35 HL135756

LJQ: R01HL111449, R01HL165718

JDC: National Library of Medicine (NLM) R01LM013154-01

Supported in part by a research grant from Drs. Margaret Seton and Joseph Jacobson in memory of Dr. Dennis J Beer.

This work was supported by the Boston University Flow Cytometry Core Facility, Microarray and Sequencing Core, and Department of Medicine Single Cell Sequencing Core. Department of Medicine Evans Junior Research Faculty Merit Award.

