## Supplemental Figures for "Neutrophil transcriptome diverges into two discrete trajectories in a murine model of severe *Streptococcus pneumoniae* pneumonia"

SUPPLEMENTAL FIGURE 1

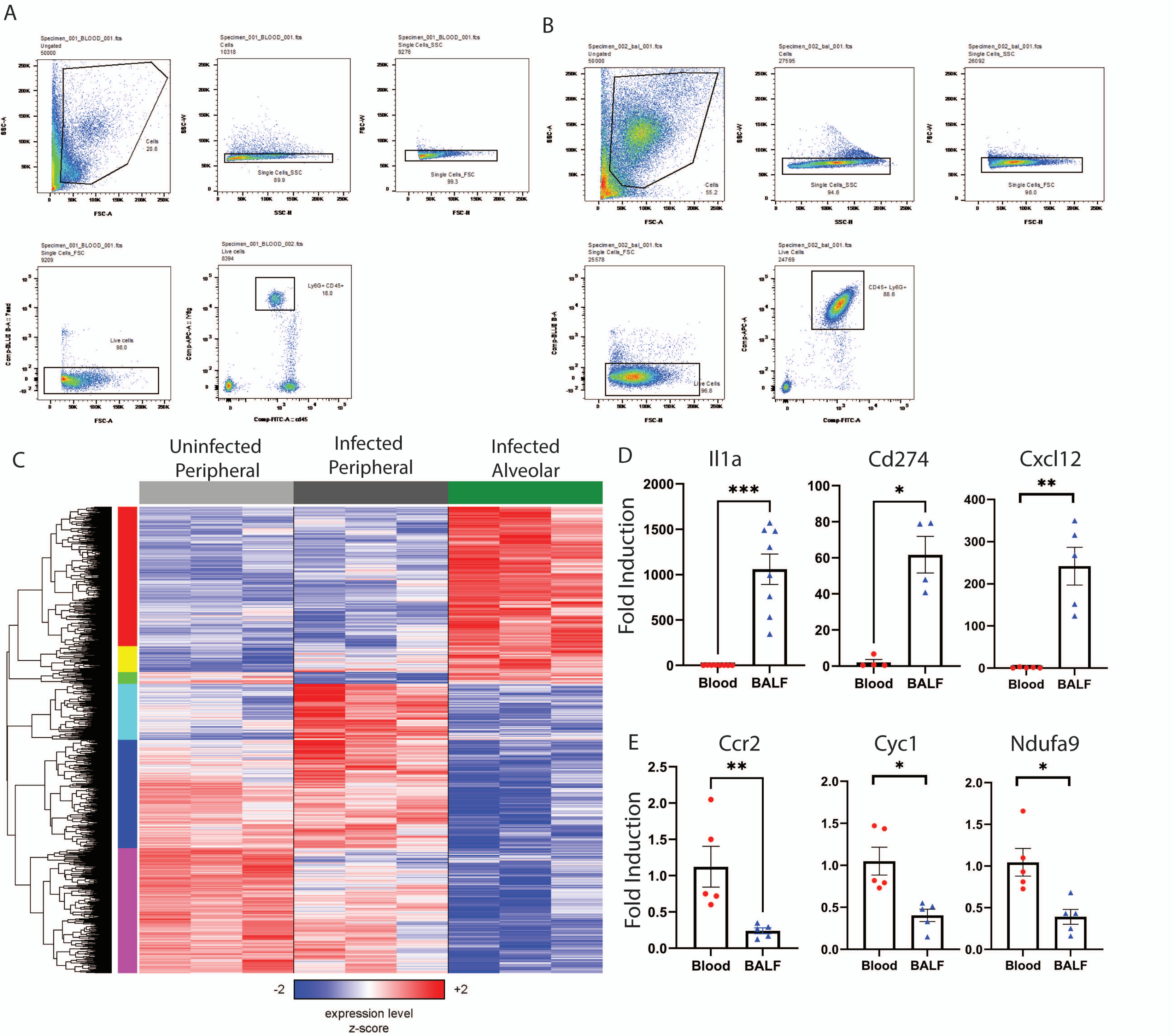


**Supplemental Figure 1.** Gating strategy is shown for isolating neutrophils by FACS sorting with examples from an infected (A) blood and (B) BAL sample. (C) Heat map of DEGs (one-way ANOVA FDR $q < 0.05$, 2783 genes), divided into six clusters by hierarchical clustering. Rows and columns correspond to genes and samples, respectively; colors are scaled by row, with blue and red indicating values that are at least 2 standard deviations below or above the mean (white), respectively. Confirmatory analysis of the blood and BAL microarray was performed by sorting blood and BAL neutrophils and performing qPCR on genes that were (D) upregulated in BALNs and (E) downregulated in BALNs. Values are fold change compared to blood neutrophil. Student's t test was performed, with P value represented by * = $P < 0.05$, ** = $P < 0.01$, and *** = $P < 0.001$.

SUPPLEMENTAL FIGURE 2

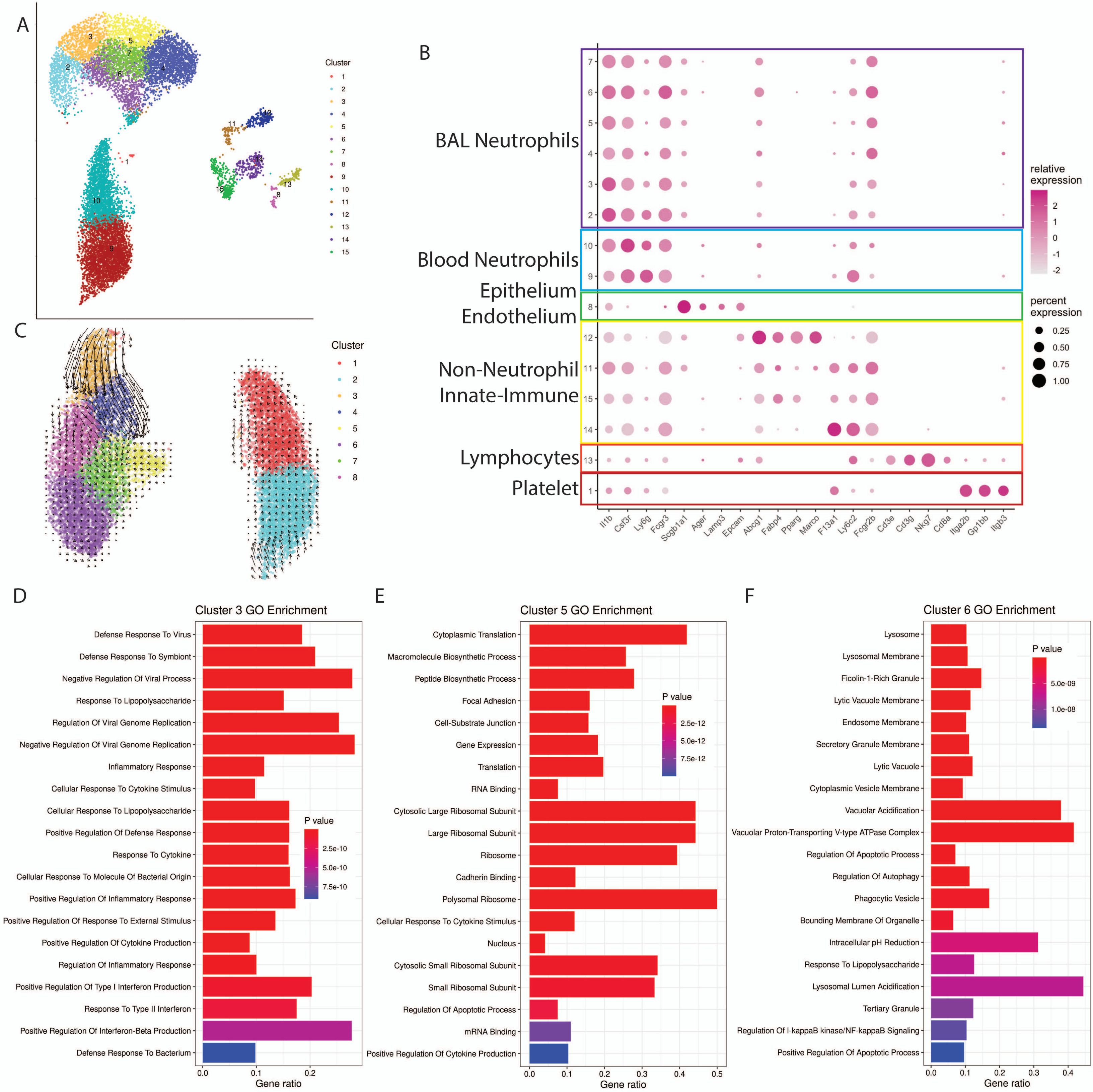


**Supplemental Figure 2.** (A) UMAP of pre-filtered scRNAseq data colored by 15 initial Celda clusters. (B) Bubble plot of markers used to identify non-neutrophil cells and clusters, with cell clusters on the Y-axis, lineage-associated genes on the X-axis and bubble size and color representing relative and percent expression respectively. (C) RNA velocity of the blood and BAL neutrophils. (D-F) Plot of top 20 GO pathways specifically upregulated in (D) early BALN C3, (E) late BALN C5, or (F) late BALN C6.

SUPPLEMENTAL FIGURE 3

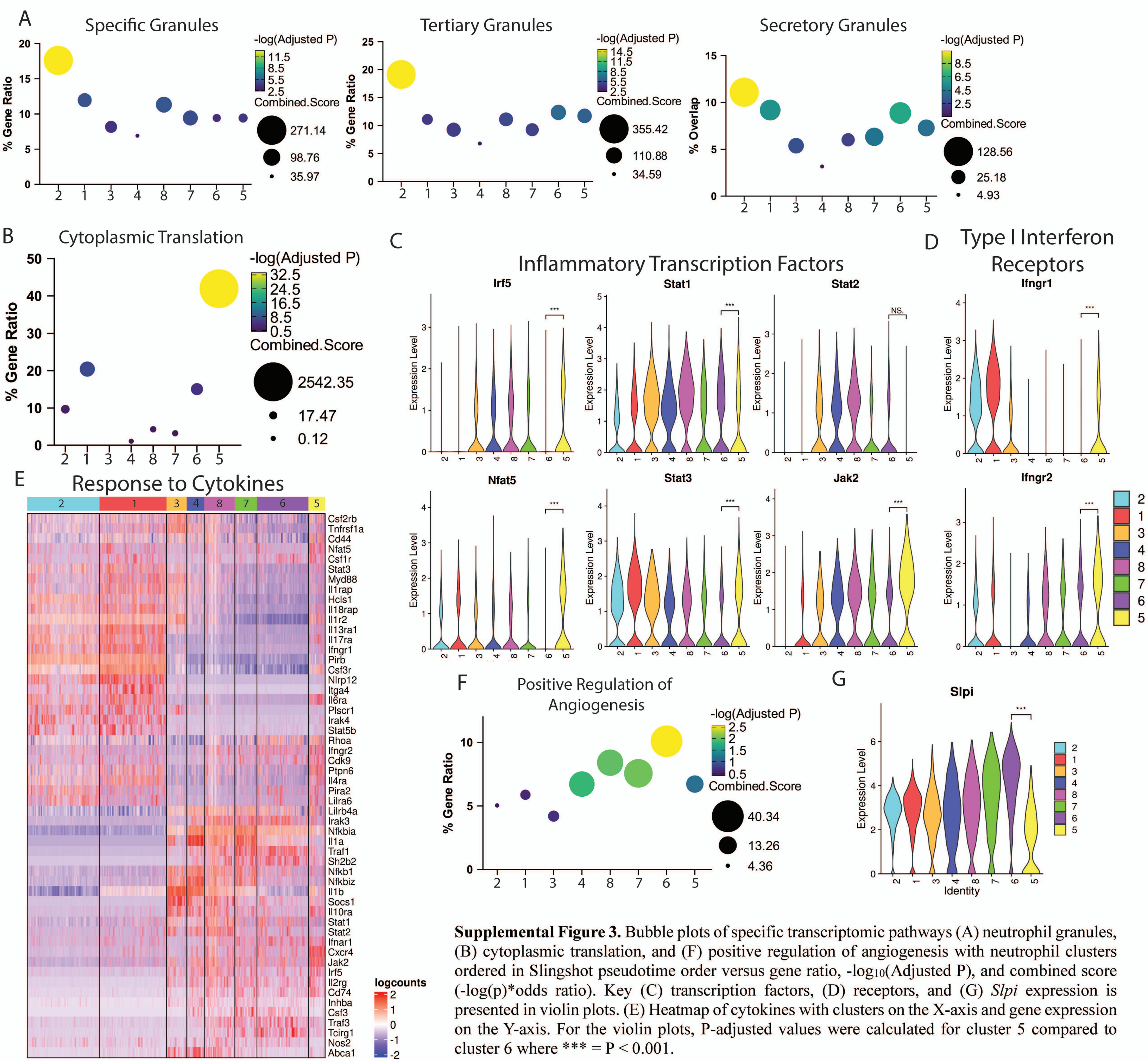


**Supplemental Figure 3.** Bubble plots of specific transcriptomic pathways (A) neutrophil granules, (B) cytoplasmic translation, and (F) positive regulation of angiogenesis with neutrophil clusters ordered in Slingshot pseudotime order versus gene ratio, -$\log_{10}$(Adjusted P), and combined score (-log(p)*odds ratio). Key (C) transcription factors, (D) receptors, and (G) *Slpi* expression is presented in violin plots. (E) Heatmap of cytokines with clusters on the X-axis and gene expression on the Y-axis. For the violin plots, P-adjusted values were calculated for cluster 5 compared to cluster 6 where *** = $P < 0.001$.

SUPPLEMENTAL FIGURE 4

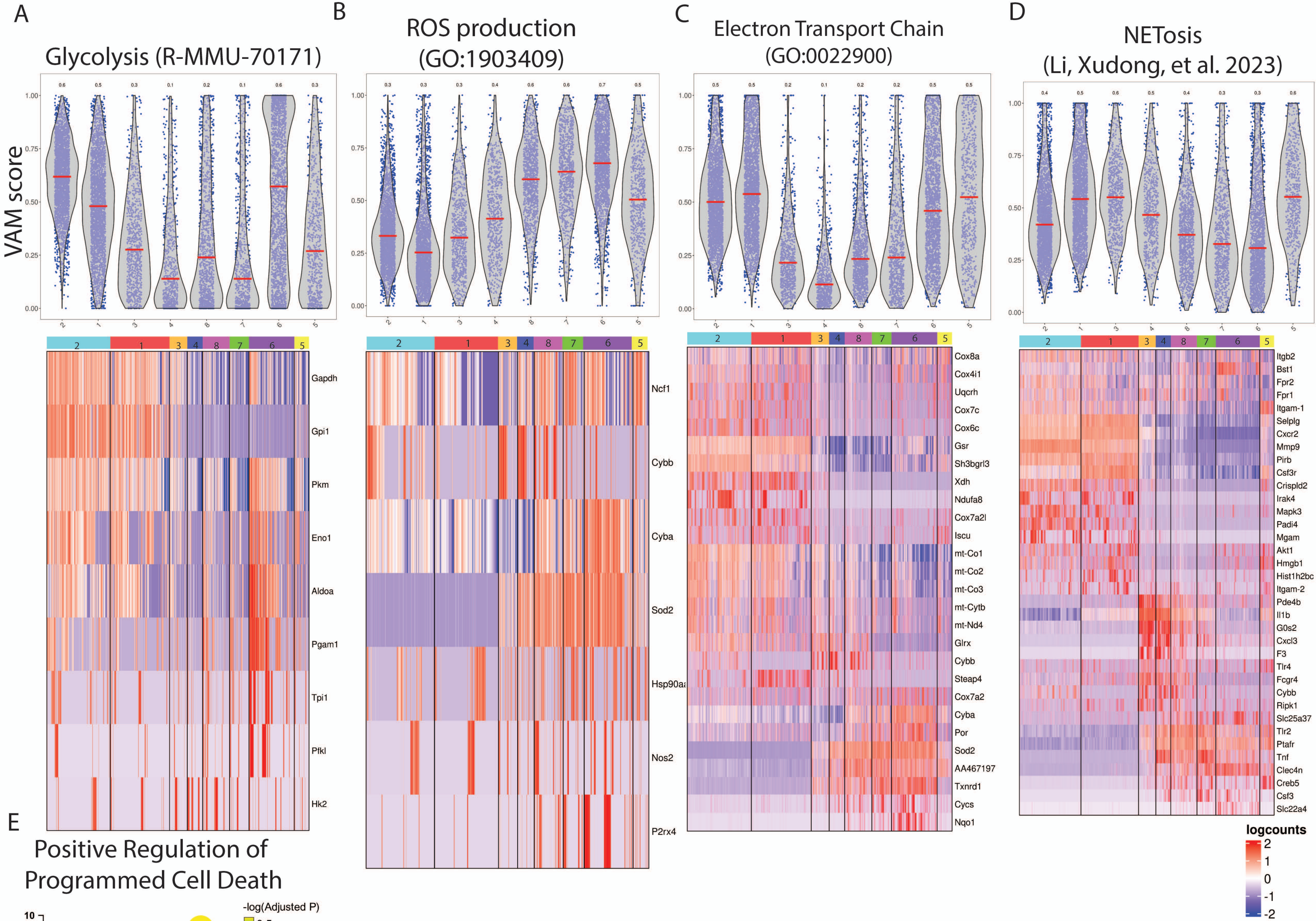


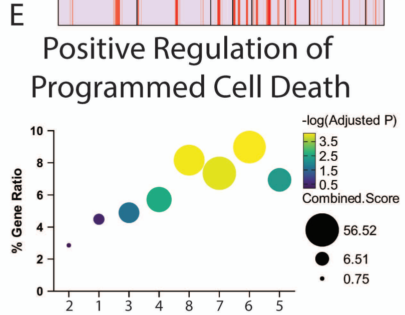


**Supplemental Figure 4.** Gene set and pathway scoring was performed via the Variance-Adjusted Mahalanobis (VAM) method. (A-D) In each pair of plots, clusters are ordered horizontally in RNA Slingshot order. The top violin plots show the gene set fitted VAM area under the curve score for each pathway. The lower plot is a heatmap of genes significantly differentially expressed from the analyzed pathway (A) glycolysis, (B) ROS production, (C) electron transport chain, and (D) NETosis. (E) The response to cell death pathway is presented in bubble plots showing the neutrophil clusters ordered in Slingshot pseudotime order versus gene ratio, $-\log_{10}$(Adjusted P), and combined score.

SUPPLEMENTAL FIGURE 5

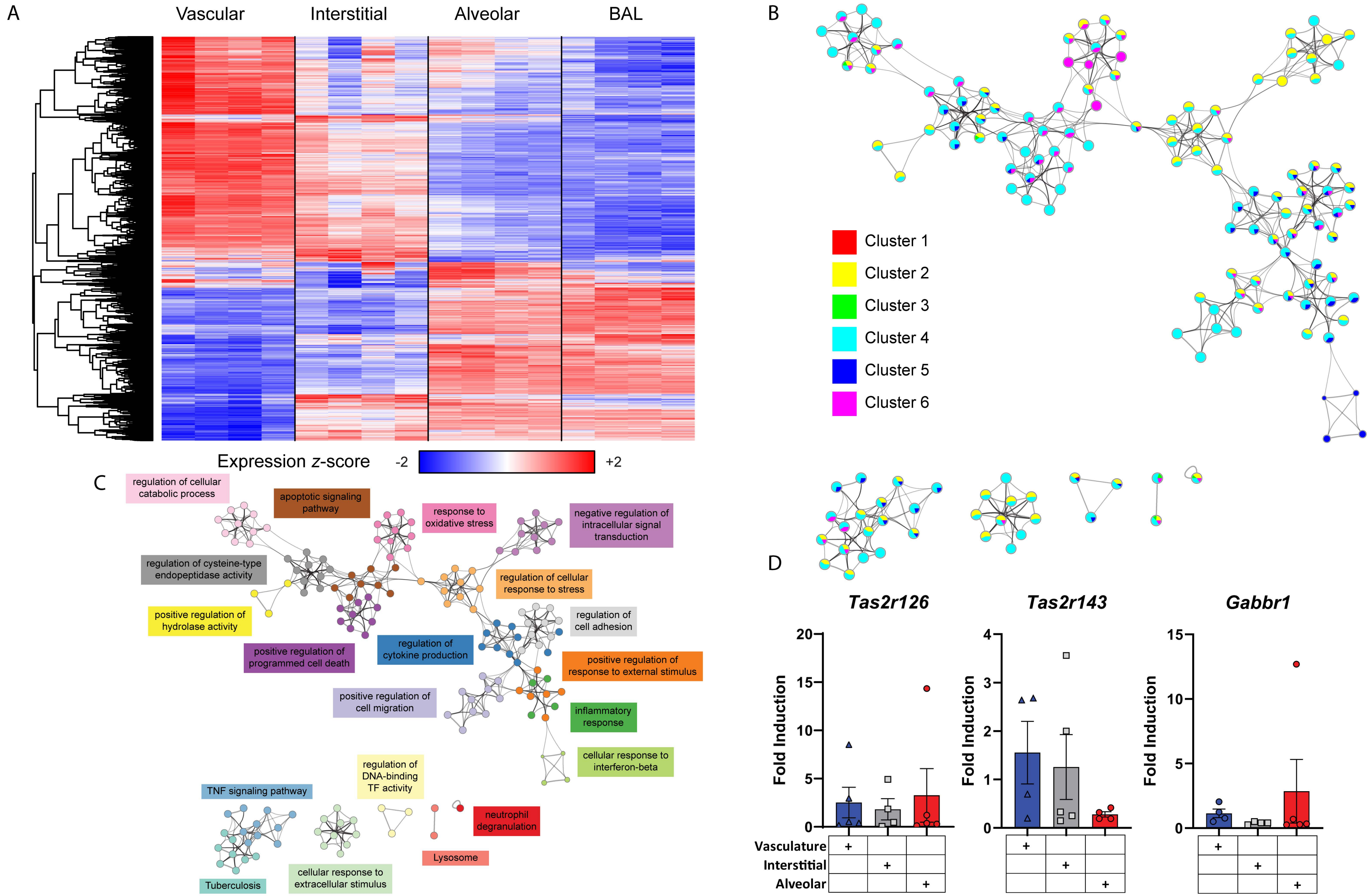


**Supplemental Figure 5.** (A) Heatmap of expression in neutrophils from all four pulmonary compartments (vascular, interstitium, alveolar airspace, and BAL). Rows correspond to DEGs (one-way ANOVA FDR $q < 0.25$, n=3924 genes), and columns correspond to samples within each compartment. Colors are scaled by row, with blue and red indicating values that are at least 2 standard deviations below (blue) or above (red) the mean (white) respectively. (B) A subset of representative terms from the Metascape analysis performed across all clusters, converted to a network layout with nodes labeled as pie charts representing the fraction of genes from each cluster. (C) The same Metascape enrichment network, in which each term is represented by a circular node whose size is proportional to the number of input genes, and terms with a similarity score > 0.3 are linked by edges whose thickness is proportional to the similarity score. Nodes are shaded by term (denoted in colored boxes). (D) Neutrophils from SP3-infected mice isolated from the pulmonary vasculature, interstitium, or airspace were stained and sorted by FACS as previously described. qRT-PCR was performed on *Tas2r126, Tas2r143,* and *Gabbr1* and normalized to expression from neutrophils from the pulmonary vasculature from uninfected mice. Differences are not statistically significant by one-way ANOVAs with Bonferroni multiple comparison corrections.
