## Supplemental Tables for "Neutrophil transcriptome diverges into two discrete trajectories in a murine model of severe *Streptococcus pneumoniae* pneumonia": 20241003 Supplemental Table 1 top 20 genes.docx

**Supplemental Table 1:** *Top upregulated (log_2_(FC)>4) DEG and top downregulated (log_2_(FC)<-2) DEG*

| **Abbreviation** | **Name** | **log_2_(FC)** | **-log_10_(FDR)** |
| --- | --- | --- | --- |
| Il1a | Interleukin 1 alpha | 7.19 | 3.15 |
| Ccl3 | Chemokine (C-C motif) ligand 3 | 6.01 | 2.78 |
| Mir155hg | Mir155 host gene (non-protein coding) | 5.49 | 2.28 |
| Olr1 | Oxidized low density lipoprotein (lectin-like) receptor 1 | 5.20 | 3.13 |
| Nr4a3 | Nuclear receptor subfamily 4, group A, member 3 | 5.17 | 2.76 |
| Cxcl10 | Chemokine (C-X-C motif) ligand 10 | 5.16 | 3.13 |
| Ralgds | Ral guanine nucleotide dissociation stimulator | 4.90 | 2.93 |
| Cxcl3 | Chemokine (C-X-C motif) ligand 3 | 4.70 | 2.11 |
| Rgs1 | Regulator of G-protein signaling 1 | 4.67 | 2.23 |
| Fam71f2 | Family with sequence similarity 71, member F2 | 4.65 | 3.13 |
| Ccrl2 | Chemokine (C-C motif) receptor-like 2 | 4.60 | 2.12 |
| Ccl4 | Chemokine (C-C motif) ligand 4 | 4.49 | 2.93 |
| Slc7a11 | Solute carrier family 7, member 11 | 4.42 | 2.90 |
| Spp1 | Secreted phosphoprotein 1 | 4.36 | 2.59 |
| Mreg | Melanoregulin | 4.36 | 2.17 |
| Il23a | Interleukin 23, alpha subunit p19 | 4.28 | 2.69 |
| Gm19510 | Predicted gene, 19510 | 4.26 | 2.81 |
| Tnf | Tumor necrosis factor | 4.22 | 2.80 |
| F3 | Coagulation factor III | 4.20 | 2.70 |
| Traf1 | TNF receptor-associated factor 1 | 4.18 | 2.27 |
| Ola1 | Obg-like atpase 1 | -2.01 | 2.10 |
| D6Wsu163e | DNA segment, Chr 6, Wayne State University 163, expressed | -2.03 | 2.18 |
| Rpl4 | Ribosomal protein L4 | -2.04 | 2.46 |
| Actr6 | ARP6 actin-related protein 6 | -2.06 | 2.08 |
| Pdlim1 | PDZ and LIM domain 1 (elfin) | -2.11 | 2.93 |
| Snx5 | Sorting nexin 5 | -2.18 | 2.11 |
| Camk2d | Calcium/calmodulin-dependent protein kinase II, delta | -2.28 | 2.13 |
| Gm10260 | Ribosomal protein S18 pseudogene | -2.32 | 2.50 |
| Pot1b | Protection of telomeres 1B | -2.53 | 2.36 |
| Crip1 | Cysteine-rich protein 1 (intestinal) | -2.62 | 3.13 |
| Scarb1 | Scavenger receptor class B, member 1 | -2.68 | 2.13 |
| Fkbp3 | FK506 binding protein 3 | -2.70 | 2.06 |
| Plekho1 | Pleckstrin homology domain containing, family O member 1 | -2.73 | 2.07 |
| Ifi30 | Interferon gamma inducible protein 30 | -2.82 | 2.12 |
| Slc25a4 | Solute carrier family 25, member 4 | -2.82 | 2.29 |
| S100a4 | S100 calcium binding protein A4 | -2.91 | 2.24 |
| Il6st | Interleukin 6 signal transducer | -2.97 | 3.01 |
| Fn1 | Fibronectin 1 | -3.21 | 2.18 |
| Adgre4 | Adhesion G protein-coupled receptor E4 | -3.48 | 2.15 |
| Pld4 | Phospholipase D family, member 4 | -4.69 | 3.13 |
