## Supplemental Tables for "Neutrophil transcriptome diverges into two discrete trajectories in a murine model of severe *Streptococcus pneumoniae* pneumonia": 20241003 Supplemental Table 2 top 20 genesets.docx

**Supplemental Table 2:** *Top Coordinately Upregulated or Downregulated Gene Sets by GSEA*

| **Gene Set Name** | **Size** | **NES** | **FDR q** | **Group** |
| --- | --- | --- | --- | --- |
| TNFα signaling via NFκB | 192 | 3.59 | <0.0001 | Hallmark |
| Inflammatory response | 190 | 3.21 | <0.0001 | Hallmark |
| Interferon gamma response | 181 | 2.97 | <0.0001 | Hallmark |
| Interferon alpha response | 86 | 2.76 | <0.0001 | Hallmark |
| Cytokine activity | 165 | 2.55 | <0.0001 | GO MF |
| Negative regulation of type I interferon production | 35 | 2.55 | <0.0001 | GO BP |
| Response to molecule of bacterial origin | 296 | 2.53 | <0.0001 | GO BP |
| Inflammatory response | 394 | 2.49 | <0.0001 | GO BP |
| Response to type I interferon | 41 | 2.47 | <0.0001 | GO BP |
| Il6 JAK STAT3 signaling | 81 | 2.46 | <0.0001 | Hallmark |
| Interferon alpha beta signaling | 39 | 2.46 | <0.0001 | RP |
| Apoptosis | 160 | 2.44 | <0.0001 | Hallmark |
| Response to bacterium | 409 | 2.40 | <0.0001 | GO BP |
| Regulation of NFκB import into nucleus | 44 | 2.40 | <0.0001 | GO BP |
| Regulation of cytokine secretion | 128 | 2.35 | 0.0003 | GO BP |
| NFκB 01 | 237 | 2.35 | 0.0003 | TF motif |
| Regulation of IκB kinase NFκB signaling | 214 | 2.34 | 0.0004 | GO BP |
| Leishmania infection | 60 | 2.33 | 0.0004 | KEGG pathway |
| Cellular response to interleukin 1 | 73 | 2.33 | 0.0004 | GO BP |
| Extrinsic apoptotic signaling pathway | 92 | 2.33 | 0.0004 | GO BP |
| Mitochondrial matrix | 392 | -2.49 | <0.0001 | GO CC |
| Translational termination | 89 | -2.49 | <0.0001 | GO BP |
| Structural constituent of ribosome | 167 | -2.50 | <0.0001 | GO MF |
| Ribonucleoprotein complex biogenesis | 383 | -2.50 | <0.0001 | GO BP |
| Ribosome | 182 | -2.51 | <0.0001 | GO CC |
| Respiratory chain | 71 | -2.51 | <0.0001 | GO CC |
| ncRNA processing | 345 | -2.52 | <0.0001 | GO BP |
| Organellar ribosome | 69 | -2.55 | <0.0001 | GO CC |
| MYC targets v1 | 185 | -2.55 | <0.0001 | Hallmark |
| Ribosome biogenesis | 269 | -2.55 | <0.0001 | GO BP |
| Ribosomal subunit | 124 | -2.57 | <0.0001 | GO CC |
| Electron transport chain | 84 | -2.57 | <0.0001 | GO BP |
| Mitochondrial protein complex | 119 | -2.58 | <0.0001 | GO CC |
| Translational elongation | 104 | -2.58 | <0.0001 | GO BP |
| Inner mitochondrial membrane protein complex | 90 | -2.58 | <0.0001 | GO CC |
| Respiratory electron transport | 57 | -2.62 | <0.0001 | RP |
| Mitochondrial translation | 104 | -2.62 | <0.0001 | GO BP |
| Respiratory electron transport ATP synthesis by chemiosmotic coupling and heat production by uncoupling proteins | 68 | -2.65 | <0.0001 | RP |
| Oxidative phosphorylation | 183 | -2.73 | <0.0001 | Hallmark |
| TCA cycle and respiratory electron transport | 103 | -2.74 | <0.0001 | RP |

NES: Normalized Enrichment Score

GO BP: Gene ontology biological process

GO CC: Gene ontology cellular component

GO MF: Gene ontology molecular function

RP: Reactome pathway
