## Supplemental Tables for "Neutrophil transcriptome diverges into two discrete trajectories in a murine model of severe *Streptococcus pneumoniae* pneumonia": 20241003 Supplementary table 7 qPCR primers.docx

Supplementary Table 4

| Target | Company | Catalog Number |
| --- | --- | --- |
| IL1a | ThermoFisher Scientific | Mm00439620_m1 |
| Cxcl2 | ThermoFisher Scientific | Mm99999117_s1 |
| Cd274 | ThermoFisher Scientific | Mm01208504_m1 |
| Cyc1 | ThermoFisher Scientific | Mm00470540_m1 |
| Ccr2 | ThermoFisher Scientific | Mm99999051_gh |
| Ndufa9 | ThermoFisher Scientific | Mm00481216_m1 |
| Tas2r126 | ThermoFisher Scientific | Mm01702063_s1 |
| Tas2r143 | ThermoFisher Scientific | Mm01700139_s1 |
| Gabr1 | ThermoFisher Scientific | Mm00444578_m1 |
| Eif2ak2 | ThermoFisher Scientific | Mm01235643_m1 |
